# Cavity geometry shapes overall ant colony organization through spatial limits but workers maintain fidelity zones

**DOI:** 10.1101/2022.06.30.498314

**Authors:** Greg T. Chism, William Nichols, Anna Dornhaus

## Abstract

Many animals inhabit nests that protect them from adverse environments. However, the effects of living in a built or found structure are not limited to protection: the physical space can shape and organize behavior, particularly in self-organized collective systems. In addition, the geometry of nest space may not be under the animal’s control, raising the question whether animals can compensate for the effects that unexpected or suboptimal geometries may have. Here we examine how the shape of a nest cavity affects spatial organization of colonies in the ant *Temnothorax rugatulus*, a species that adapted to nest cavities of unmodifiable internal dimensions, since they inhabit rock crevices with rigid walls. We show that the emerging spatial relationships of workers, brood, queens, and young alates, as well as their relationships and distances to significant points in the nest, are all significantly influenced by nest shape, with the brood distributions most affected. However, we also found that the size of worker spatial fidelity zones, i.e. the areas in the nest that individual workers occupy and that may be key regulators of division of labor, are overall not affected by nest shape. These findings indicate that ants may actively regulate which areas of a nest they occupy, and that they may compensate for effects of nest architecture constraints. Physical properties of nests can thus influence the organization of ant colonies, highlighting the need to explore spatial constraints as a direct influence on the organization, movement, and communication of evolved or engineered self-organized systems.

## Introduction

Social animal architectures benefit individuals by opening a wider range of suitable habitats that they, and their kin, can successfully inhabit. These architectures, i.e., built environmental modifications, range from simple depressions in the ground (e.g., social crab burrows, Laidre et al., 2018) to massive above and below ground structures that span meters across (e.g., *Macrotermes* termites, Harris 1956). This diversity is unified by the impact each architecture can have on the social behavior of their occupants: social weaver bird nests promote altruism through spatial and social clustering (van Dijk et al., 2015), kin cohabitation increases with burrow size in social crabs (Laidre et al., 2018, Laidre 2019), and reproductive division of labor is facilitated by both naked mole rat burrows (Tofts and Franks 1992; Faulkes and Bennett 2001) and social insects nests (Wilson and Kinne 1990, Wilson 1992). Ant nests are particularly interesting because of both their representation in nearly every environment, and their diversity in form and function within these environments (Wilson and Kinne 1990, Wilson 1992).

Here we are interested in how the geometric shape of such built architectures influences the organization of the colony within. Ants have been demonstrated to be distributed non-uniformly and non-randomly across their internal nest space in many cases (Sendova-Franks and Franks 1995; Sendova-Franks and Franks 1999; Tschinkel 1999; Tschinkel 2005). For example, colony members may be concentrated in specific sections of the nest (Tschinkel 1999; Tschinkel 2005); and workers may use chemical ‘road-signs’ to navigate nest space and differentiate chambers with different functions (Heyman et al., 2017). However, some complex spatial relationships in the nest may be predicted by random walk models (Sendova-Franks and Van Lent 2002, Davidson and Gordon 2017), implying that the placement of colony members may not always be an explicitly adaptive strategy. In addition, many ants exploit pre-existing natural structures as nests (e.g., *Temnothorax* ants: Prebus 2017, *Cephalotes* turtle ants that occupy unmodifiable wood nests: Powell 2008, and rock dwelling *Rhytidoponera metallica* ponerine ants: Thomas 2002), which means that the geometry of the available space may not be under the ants’ control. We therefore ask how the internal nest space may influence the spatial organization of workers, queens, and brood in the nest.

It has become clear over the last decade that the pattern of interactions in and allowed by social insect nests may affect colony organization. For example, worker interactions and recruitment increase in nest chambers that are highly connected to other nest chambers (Pinter-Wollman 2015; Vaes et al., 2020), information processing and exploitation of the environment decreases in nests with multiple entrances (Lehue and Detrain 2019; Lehue and Detain 2020; Lehue et al., 2020a,b; Lehue and Detrain 2020), while panicked workers bottleneck when evacuating through a single nest entrance (Burd et al., 2010; Wang and Song 2016). Further, worker density in a nest increases the likelihood that colonies become polydomous in the ant *Temnothorax rugatulus* (Cao 2013). Social interactions and heterogeneity in spatial distributions in the nest are closely linked (Guo et al., 2020). Information, such as alarm signals, may be spread through worker physical contact, in which case the spatial arrangement of workers critically affects information flow (Guo et al., 2022). Further, queen(s) and brood may be positioned in nests so that they are both more accessible from other nest spaces and possibly more defensible during nest invasions (Varoudis et al., 2018). Ant colonies further reduce disease spread through social distancing: disease spread is reduced by adaptively clustered (rather than globally random) worker interactions, and patterns of interactions thus may confer organizational immunity (Stroeymeyt et al., 2018; Pusceddu et al., 2021; Naug and Smith 2007). It thus seems clear that worker (and other colony member) interactions and density may affect colony-level performance in essential tasks both inside and outside the nest; and thus, properties of the inhabited space, which are likely to affect such patterns of interaction and spatial distribution, are in turn critical to understand collective colony function.

One context in which spatial relationships appear particularly important is task allocation and division of labor. Although this may vary between tasks and species, proximity of individuals to areas where particular work is needed seems to be a major factor affecting task allocation (Leitner and Dornhaus 2019). Workers do not move through the nest randomly, and are often faithful to particular, often quite small, mini-‘territories’ in the nest called ‘spatial fidelity zones’. Spatial fidelity zones have been demonstrated in multiple species, and the location of these zones often correlates with worker task (Franks and Tofts 1994; Sendova-Franks and Franks 1994; Sendova-Franks and Franks 1995; Sendova-Franks and Franks 1999; Powell and Tschinkel 1999; Jandt and Dornhaus 2009). The size of these fidelity zones varies with task, location in the nest, or specialization of workers (e.g. smaller fidelity zones in workers performing brood care, Sendova-Franks and Franks 1995; Jandt and Dornhaus 2009, or inactive workers, Charbonneau et al., 2017a, Leitner and Dornhaus 2019).

Nest-related collective behavior has been well studied in the ant genus *Temnothorax*. Rock-dwelling *Temnothorax* have been the subject of studies involving house-hunting (Franks et al., 2002; Dornhaus et al., 2004; Sasaki and Pratt 2013; Sasaki et al., 2015), collective nest building (Franks et al., 1992; Franks and Deneubourg 1997; Aleksiev et al., 2007a; Aleksiev et al., 2007b; Aleksiev et al., 2007c; DiRienzo and Dornhaus 2017), and risk-taking behaviors relating to ecological nest site availability (Bengston and Dornhaus 2014; Bengston and Dornhaus 2015; Bengston et al., 2017), among other topics. We used the rock ant *Temnothorax rugatulus* to examine the influence of nest shape on its occupants. *T. rugatulus* colonies dwell in pre-existing rock-crevices, producing single-chambered nests that are easily replicated and manipulated in a laboratory setting (Charbonneau and Dornhaus 2015). Worker spatial position in *Temnothorax unifasciatus* (a con-gener to *T. rugatulus* with similar rock-dwelling nesting behavior) nests directly influences the task they perform, such as brood care near the brood pile (Sendova-Franks and Franks 1995), and *T. unifasciatus* workers will return to their spatial positions in relation to one another in new nests (Sendova-Franks and Franks 1994). This species thus provides a suitable model system, in which we know that worker interactions and spatial arrangement are relevant to the colony, but also are feasible to comprehensively monitor and manipulate; and furthermore, ants in this genus likely have to adapt frequently to nest geometries that are largely beyond their control.

We investigate the influence of internal nest geometry on colony organization in *T. rugatulus* colonies for two artificial nest shapes designed to differ markedly in physical accessibility: a ‘circle’ and a ‘tube’ nest (Fig. 1). We quantify the distribution of all types of individuals in the nests, and then ask how the location of the nest entrance and the centroid of the brood distribution relate to where colony members are located in the nest. We also examine how the distributions are affected by overall colony density in the nest. Finally, we determined whether the spatial fidelity zones of individual workers are different across nest shapes, and how they relate to the location of nest and colony features.

**Figure 1.**
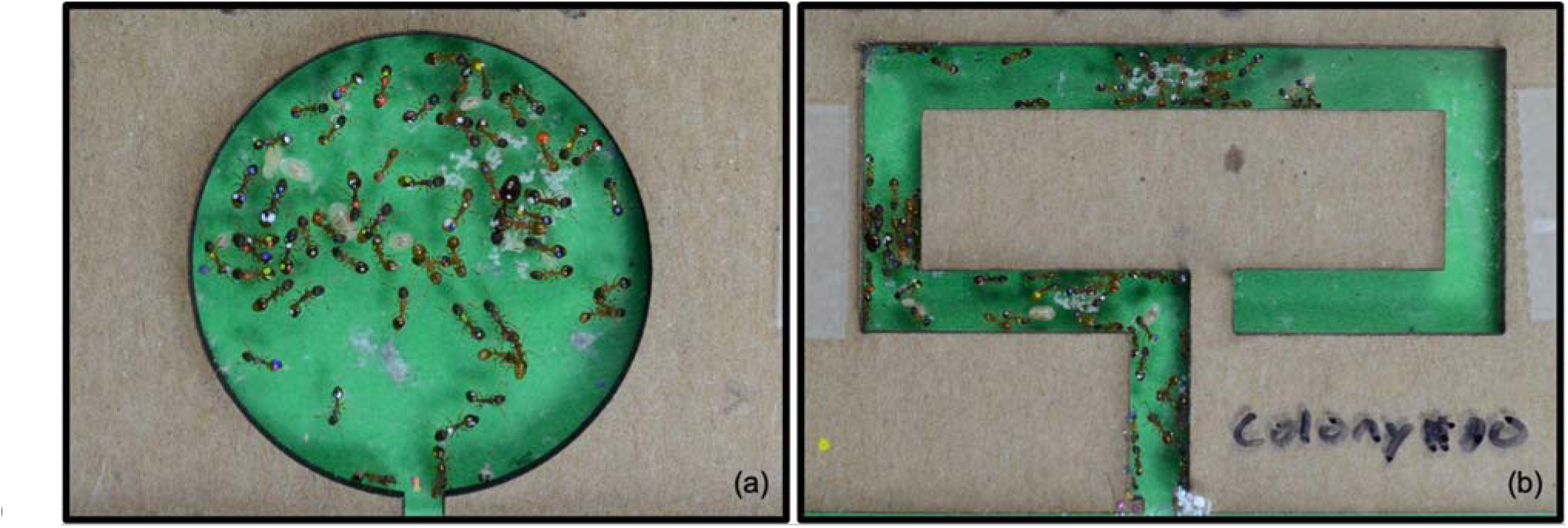
Nest shapes used for this study: (a) circle and (b) tube nests. Internal nest area was always scaled to the number of workers in the colony. The nest entrance is in the middle lower edge of the image in both cases. A green background was used to make the brood items (white) more visible. Pictured here are high-density treatment nest shapes for one colony.

## Methods

### Colony collections

We collected 20 queenright colonies of *Temnothorax rugatulus* from rocky semi-steep slopes on the Santa Catalina Mountains (GPS: 32.395, −110.688), USA, Pima County, Arizona from July to October 2017 and February to May 2018. We found all of our colonies in a pine-juniper zone (altitude approximately 2500m) inhabiting granite rock-crevices where entire colonies were collected by aspiration after prying open the nest cavity. The second half of the colonies (IDs 11-20) were collected during the colonies’ reproductive period, which allowed us to examine alate spatial organization (alates being winged ants, i.e. new queens and males). The first ten colonies had only workers and (non-alate, i.e. reproductive) queens and brood, whereas seven of the latter ten colonies additionally had alates. We did not identify or account for the presence of microgynes (diminutive but possibly reproductive queens) in our study and only identified obvious queens (significant size and color difference compared to workers and proximity to a cluster of her brood). Microgynes were likely present however, given that they have been shown to be prevalent in higher elevation populations of this species (Choppin et al., 2021). Microgynes were most likely categorized as workers in our study. They are likely to be few in number if they occur and behave and are treated differently than macrogynes (regular, large-size reproductive queens) by workers (fed more but groomed less, Negroni et al., 2021).

### Initial housing and care

We initially placed all colonies in generic artificial nests resembling their natural nest sites in rock crevices. These nests consisted of a 2-mm-thick piece of cardboard (75 mm x 50 mm x 2 mm) sandwiched between two glass panes (76.2 mm x 50.8 mm x 0.5 mm), with a 2 mm x 2 mm entrance at the center of one long side leading to an open nest space (35 mm x 25 mm) (Charbonneau et al., 2015; Fig. A1). We gave colonies food and water *ad libitum*, refreshed weekly, for the duration of their housing: water was given through water-filled, cotton-ball-stopped 5 ml plastic tubes, and food was given through 2 ml microcentrifuge tubes of honey water with a concentration of 1/4 teaspoon honey per 50ml water, and 1/8 (approximately 0.075 g) of a fresh-frozen cockroach (*Nauphoeta cinerea*). We kept colonies on a 12:12 hour light cycle (8 a.m. to 8 p.m.), constant temperature (approximately 21-24 °C). We placed all artificial nests in open-top plastic containers (11.1 cm x 11.1 cm x 3.3 cm) with walls lined with ‘insect-a-slip’ (BioQuip 2871A, ‘fluon’) to prevent escape. We individually marked CO_2_-anesthetized workers with four identifying marks with multicolored paints (Testor’s Pactra® paint): one each on the head and prosoma, and two on the gaster (Charbonneau and Dornhaus 2015). We marked ants three to five days before the colony was placed into experimental nests.

### Experimental setup

We used two nest types, a circle and a tube shape laser-cut from the 2mm thick cardboard, for the experimental phase (Fig. 1). We gave colonies comparable nest densities by scaling the internal area of the nests to colony size (number of workers). We produced nests with high worker density, by examining the size of a *Temnothorax rugatulus* colony that utilized nearly all the available nest space in a pre-experimental nest (248 workers, Fig. A1) and doubling the area per worker to permit flexible nest space use (0.033 mm^2^ per worker). We further produced a low nest density treatment by doubling the nest area allocated to each worker compared to that first treatment, producing a density half as dense as the high-density treatment (0.066 mm^2^ per worker). It is known that colonies favor nests that allow low in-nest worker densities (Franks et al., 2006; Visscher 2007; Mitrus 2015). However, these populations are thought to be nest-site-limited (Foitzik et al., 2004; Bengston and Dornhaus 2015), such that under conditions of high nest competition, colonies may often nest in sub-optimal nests or be prevented from expanding into multiple nests despite high density.

### Experimental timeline

Each colony experienced both nest shapes and both density treatments. We first randomly assigned each colony one of the nest shapes, such that ten of our colonies experienced the circle nest first and assigned the other ten colonies the tube nest first. We then forced an emigration into the new assigned nest by offering that nest and removing the top glass pane from their current nest, which prompts the ants to emigrate into the new cavity. After this emigration, we allowed the ants to acclimate to the new nest for three days, and then photographed colonies in this nest each day for 16 days. Once the ‘nest assignment I’ phase of 16 days was completed, we forced an emigration into the other nest type, and repeated the three-day acclimation and 16-day monitoring phase (‘nest assignment II’). Photos were taken with a digital HD SLR camera (Nikon D7000 with 60 mm lens). We used the first 10 photos from each nest assignment for analyses excluding only images that were problematic (e.g. out of focus).

### Photo analysis

We assigned cartesian coordinates to each colony member on the digital images and standardized (i.e. translated them back into real distances in mm by using the bottom-left and top-right outside corners of the nest as reference coordinates) them with the image analysis software *Fiji* (Schindelin et al., 2012).

### Nest sections and colony member densities

We divided each nest into eight equal-area sections from the nest entrance to the back of the nest (1-8, see Fig. A2a-b). We chose not to use more than eight bins because these nest section assignments were likely large enough to capture and segregate different worker tasks. We determined all colony member densities by calculating what proportion of total observed individuals of that type on the same image were found in that section.

### Scaled distances in the nest

#### Distance to the nest entrance

We calculated the shortest linear distance *within the available nest cavity* from each colony member to the nest entrance. In the circle nest, we straightforwardly calculated each colony member’s distance to the nest entrance (per a scaled reference x and y coordinate) using the formula: sqrt((colony member x - entrance x)^2^ + (colony member y - entrance y)^2^), with a small adjustment for the nest entrance tunnel. In the tube nest however, a direct (i.e., straight-line) path would not reflect the actual walking distance. Thus, we calculated the distance as the shortest possible line within the nest cavity to the entrance (see Fig. A3 in supplementary information for details and illustration).

#### Distance to the brood center

We calculated the brood center as the average brood x and y coordinates in the nest section with the most brood items in each observation. This was done because calculating the overall centroid of brood across the entire nest would often take this ‘center’ outside of the nest space boundaries in the tube nest. We then found the distance from each mobile colony member to this brood center (similar to the calculation for distance to the nest entrance, this was calculated as straight-line distance in circle nests, and as walking distance in the tube nest; for details see Fig. A3 in supplementary information). *Distance scaling*: Actual possible distances varied across nests since the nest dimensions were scaled to colony size to standardize worker density across colonies (see above). To be able to compare relative distributions across colonies, we scaled all calculated distances to nest size, by setting the shortest distance from the back of the tube nest shape to the entrance as 1. These ‘scaled’ distances are thus only relative to the size of that colony’s tube nest, not relative to each nest shape. Since the farthest point from the entrance in the circle nest is never as far as the farthest point in an equal-area tube nest, the maximum ‘scaled’ distance in the circle nest for any colony is thus 0.34 (i.e. 34% of the maximum available distance in the tube nest).

### Site fidelity

We generated twenty-four total zones by again dividing each of the eight nest sections described above into three (illustrated in Fig. A2). We assigned each observation of each worker with identifiable unique color markings to one of these zones. We defined ‘occurrence zone size’ for each worker as the total number of nest zones that the worker was observed in (out of the 24 total possible zones). We defined ‘spatial fidelity zones’ as the number of zones in which the worker was observed for at least 15% of its observations (note that zones, as nest sections, are all equal area for the same colony but reflect different amount of area across colonies because the total nest size is scaled to colony size). We also analyzed the true area (in cm^2^) of site fidelity zones (by adding the areas of the respective nest zones) in supplemental materials. We used this method instead of, for example, a minimum convex polygon (Jandt and Dornhaus 2009; Charbonneau, et al., 2017a) to ensure that we did not include areas that were not part of the nest cavity in the ‘tube’-shaped nest. We only included workers that were observed at least seven (out of ten possible) times (136 out of 941 possible workers in the high-density treatment and 247 out of 838 possible workers from the low-density treatment). This loss occurred in part because although all workers were initially individually painted, many lost two or more of their four color-combination marks.

While the 7 observations cutoff and the 15% threshold are both somewhat arbitrary, we reasoned that they provide a sensible combination given the total number of observations available. For a worker with between 7 and 10 observations, every zone it is observed in at least twice will be counted towards its spatial fidelity zone (2/7=29%), but zones in which it is observed only once will not be counted (1/7=14%). This method of calculating spatial fidelity thus enables us to identify zones the worker is likely to return to and constitutes a compromise between excluding too many observations/workers and reliability of the calculated zone sizes.

### Data processing

We conducted all data processing using the statistical software R (*v4.3.1*; R Core Team 2017) and RStudio (*v2023.06.1+524*; Allaire 2012), specifically the Tidyverse language (‘tidyverse’ *v1.3.1*: Wickham et al., 2019). All original data and analysis scripts are publicly available on GitHub at https://github.com/Gchism94/NestArchOrg (Chism 2022).

Colony distributions: We used linear mixed effects models to examine colony member distributions (proportion in each nest section) in our experimental nests using the R package ‘lme4’ (*v1.1-27.1*; Bates et al., 2014), where *P* values were calculated through the R package ‘lmerTest’ (*v3.1-3*; Kuznetsova et al., 2017). We report the relationships between colony member distributions across nest sections (quadratic term) and nest shape though the interactions term (nest * nest section^2^), and report how nest density affects this relationship through a three-way interaction term (Nest * Nest section^2^ * Nest density). Our linear mixed effects models here and below (unless noted otherwise) had colony identification (hereby termed ‘Colony ID’) as a random effect, and by comparing the variation explained by the fixed effects alone (marginal *R*^2^) and with the random effect included (conditional *R*^2^), we determined the amount of variation that Colony ID explained (marginal and conditional *R*^2^ values calculated through the R package ‘sjPlot’ *v2.8.14*; Lüdecke 2023).

Scaled distances: We used linear mixed effects models to examine the relationships between nest shape and individual scaled distances from the nest entrance and the center of the brood, including the interaction of nest shape and density (Nest * Nest density).

Worker site fidelity: We used linear mixed effects models to test the effect of nest shape, density, colony size, and colony ID on worker spatial fidelity and occurrence zone sizes. We also consider the true zone sizes (in cm^2^).

Worker site fidelity and distances in the nest: We used the same linear mixed effects models as above to test the effect of either worker mean scaled distance to the nest entrance and nest shape, and a separate linear mixed effects models to test the effect of worker mean scaled distance to the brood center and nest shape on site fidelity. We additionally tested the interaction terms between nest shape and colony size, nest shape and mean scaled distance to the nest entrance, and colony size and nest density.

## Results

### Overall colony distributions in the two nest shapes

All types of colony members’ distributions were significantly affected by nest shape.

Most prominently, brood typically formed a distinct, dense, oval cluster in the circular nest, often in the center or towards the back of the nest. In contrast, in the tube nest, brood was typically found across the entire length of the nest and arranged in several smaller clusters, which led to more brood (and workers) being located nearer the entrance of the nest (Fig. 2). We analyzed both how ants were distributed in the nest sections, and thus how they were divided up across the available space in the nest (details in Table 1), and the distribution of their actual distances from the nest entrance (details in Table A1) and center of the brood (understood to reflect the meaningful center of the colony; details in Table A2). Broadly, nest shape affected all colony member distributions across nest sections except that of queens (Fig. 3: LME; n=3192 proportions in nest sections for workers, corresponding to 10 colonies over 4 treatments with 8 sections and 10 replicates each; similar numbers for brood and queens, n=456 for alates; all p<0.01 for nest shape as a main effect, except for queens p=0.12). Density overall had no main effect on distribution across nest sections, except for brood (p<0.03; Fig. 3), and ‘Day’, i.e. time since emigration into the nest, also had no effects (all p>0.05), indicating that colonies settled into their distributions by the beginning of data collection (3 days since emigration). Colony ID, as the only random effect included in our model, did not explain any variation (identical marginal and conditional R^2^ values, Table 1).

**Figure 2.**
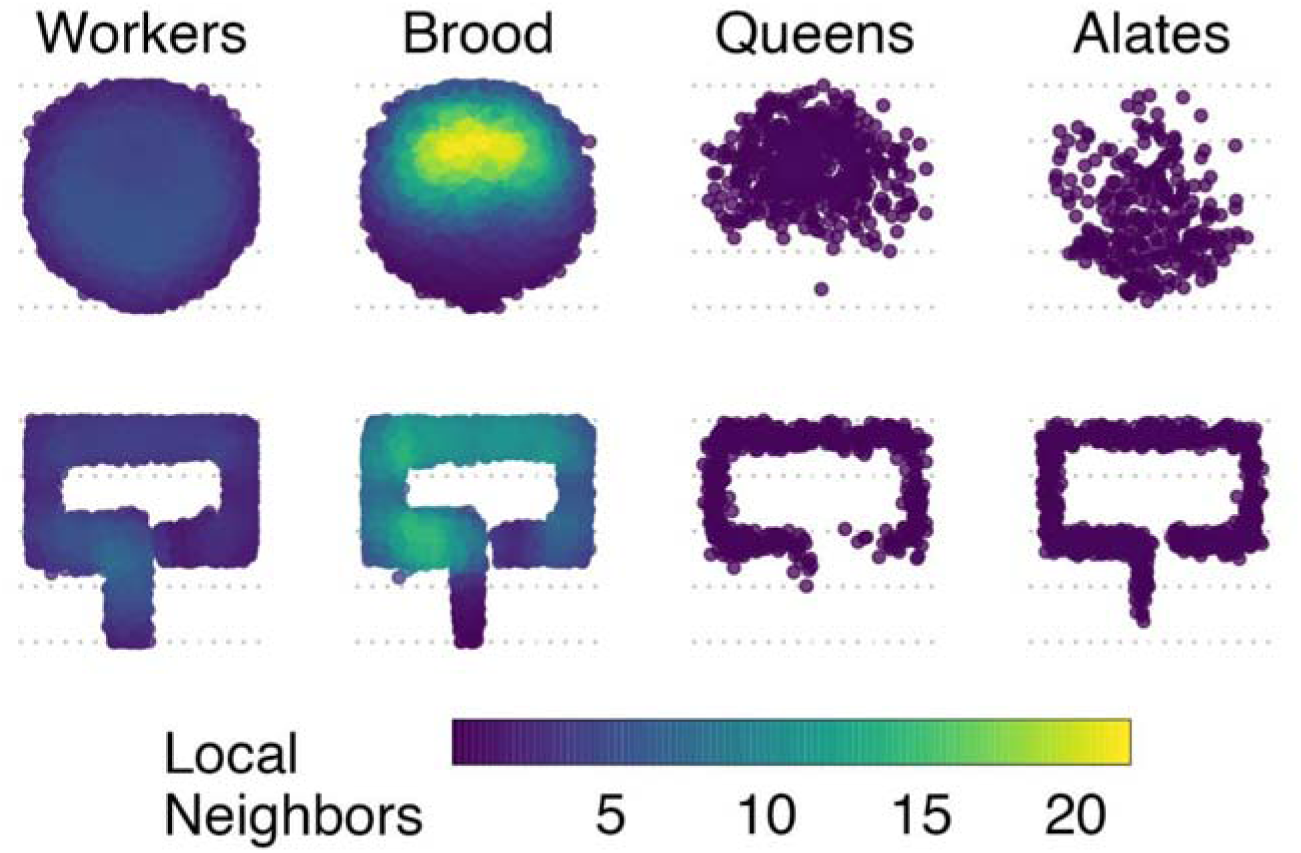
The distribution of all types of ants in the two nest shapes (top row: circle nest, bottom row: tube nest, pooled across all 400 photos from 20 colonies). Colors indicate the average number of neighbors for each item within 0.16 of the longest distance in tube nest.

**Figure 3.**
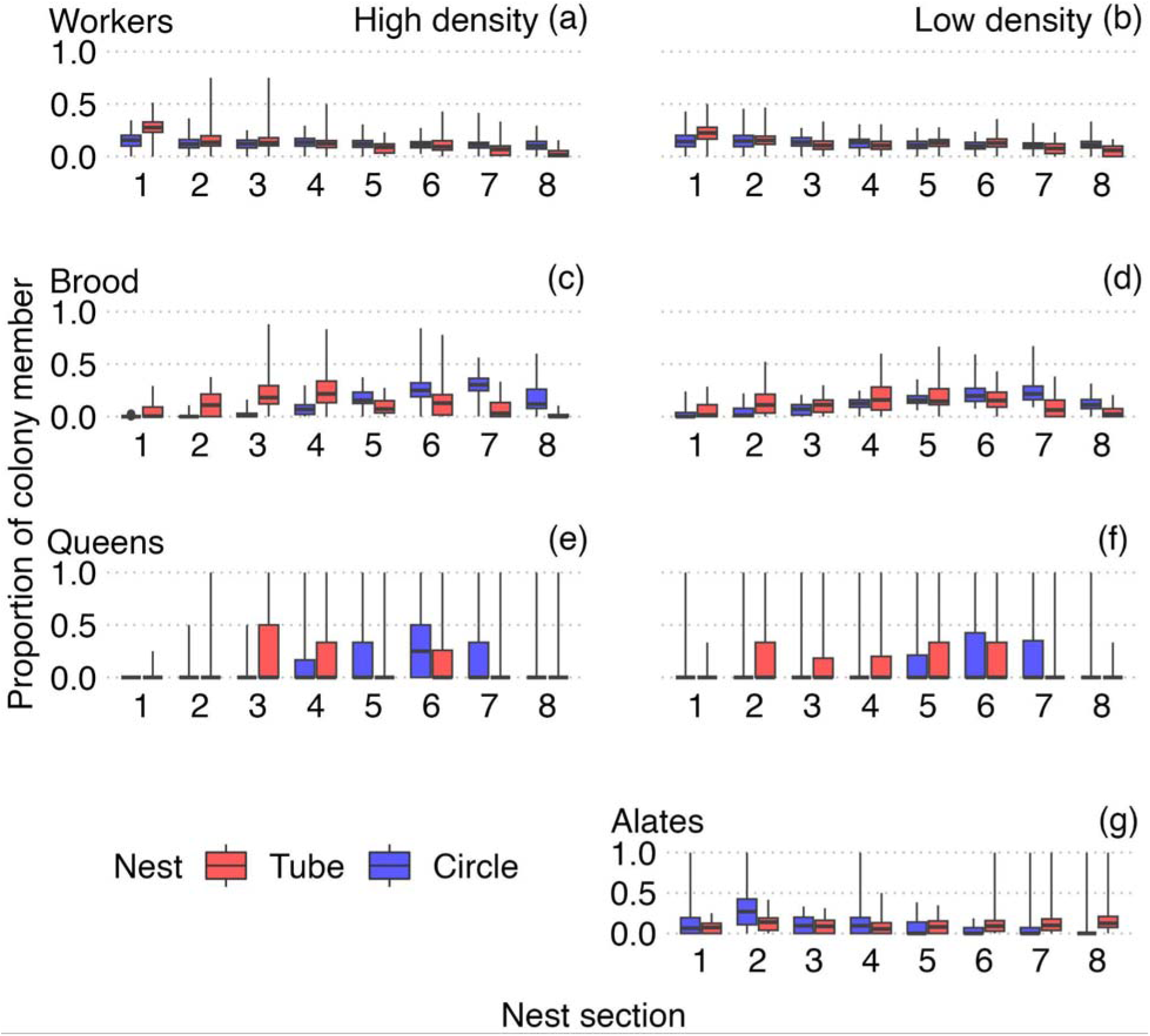
While workers are fairly evenly distributed across all available nest sections, brood and queens are clustered in intermediate areas in both nest shapes. Shown here are the proportion of the respective ant types across the equal-area nest sections: workers (a, b), brood (c, d), queens (e, f), and alates (g) in the tube (red) and circle (blue) nests and when at low (right column) or high density (left column). Nest sections scale with colony size, worker density is kept constant. Boxes represent first and third quartiles, the bar within boxes is the median, and whiskers show the data range. For corresponding statistics see Table 1.

**Table 1.**
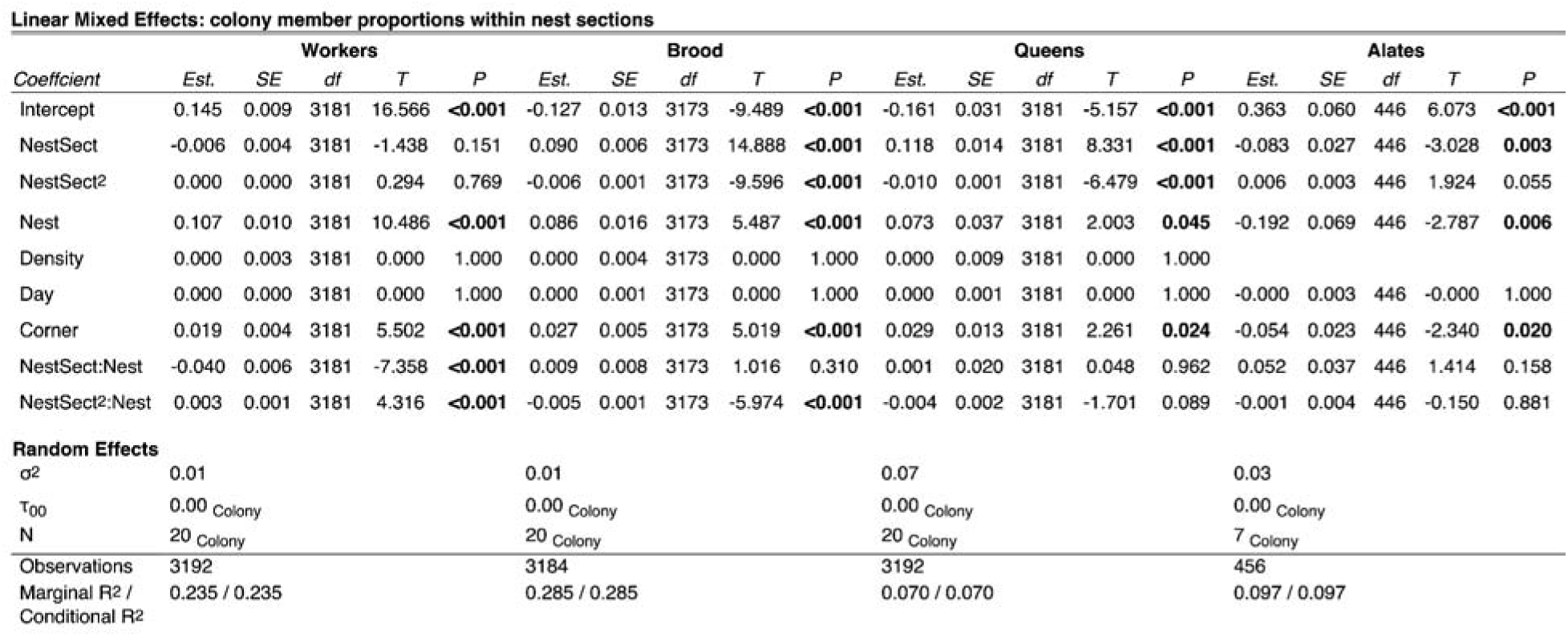
Proportions of colony members in each nest section related to nest shape and physical properties. Here and below sections (NestSect) are numbered beginning at the nest entrance (1) and end at the back of the nest (8); a quadratic term for nest section is included to allow for a unimodal distribution shape. Density is the amount of area allocated for each worker (high = 0.033144 cm^2^, low = 0.066288 cm^2^). Day is the observation day (starting 3 days after colonies moved into the nest). Corner is the presence of a bend in the tube nest or the corners around the entrance tunnel in the circle nest in the respective nest section. The random effect Colony ID is colony identity. Bold *P* values indicate significance. Model formula in R: PropColonyMember ∼ poly(NestSect, degree = 2, raw = TRUE)*Nest + Density + Day + Corner + (1 | Colony ID)

### Distances from the nest entrance

For both brood and queens, there were significant linear and quadratic effects of ‘nest section’ (all p<0.001), indicating that both distributed with an intermediate peak along the axis from the front to the back of the nest, whereas there were no main effects (linear or quadratic) of nest section on worker distributions (p=0.85 and p=0.71), indicating workers were largely evenly distributed across the nest space (as can also be seen in Fig. 2 and 3). However, nest shape interacted significantly with nest section for worker distributions, and with the quadratic effect of nest section for brood - indicating a change in the shape of the worker and brood distributions across the extent of the nest with nest geometry.

When analyzing the specific, individual, scaled distances of each ant from the nest entrance (Fig. 4), we found that nest shape again influenced all colony members’ distributions (including that of queens: LME, n=30247 locations of workers, 59459 locations of brood items, 1041 locations of queens, 1006 locations of alates, all p<0.001, Table A1): all were on average further from the nest entrance in the tube nest. This result is not very surprising since the tube nest allows ants to be located much further from the entrance than space is available in the circle nest. However, it illustrates that while there are also effects of relative spread over the nest as stated in the last section, there are obvious effects on real distances between ants when nest geometry allows those to be larger (illustrated in Fig. 4). We also found that worker density in the nest did not affect distances to the entrance (of any ant types, all p>0.05), and male alates were closer than queen alates to the nest entrance (p<0.001 for the main effect of sex for alates). Over time, workers and brood tended to move further, and alates closer to the nest entrance (effect of Day, all p<0.001).

**Figure 4.**
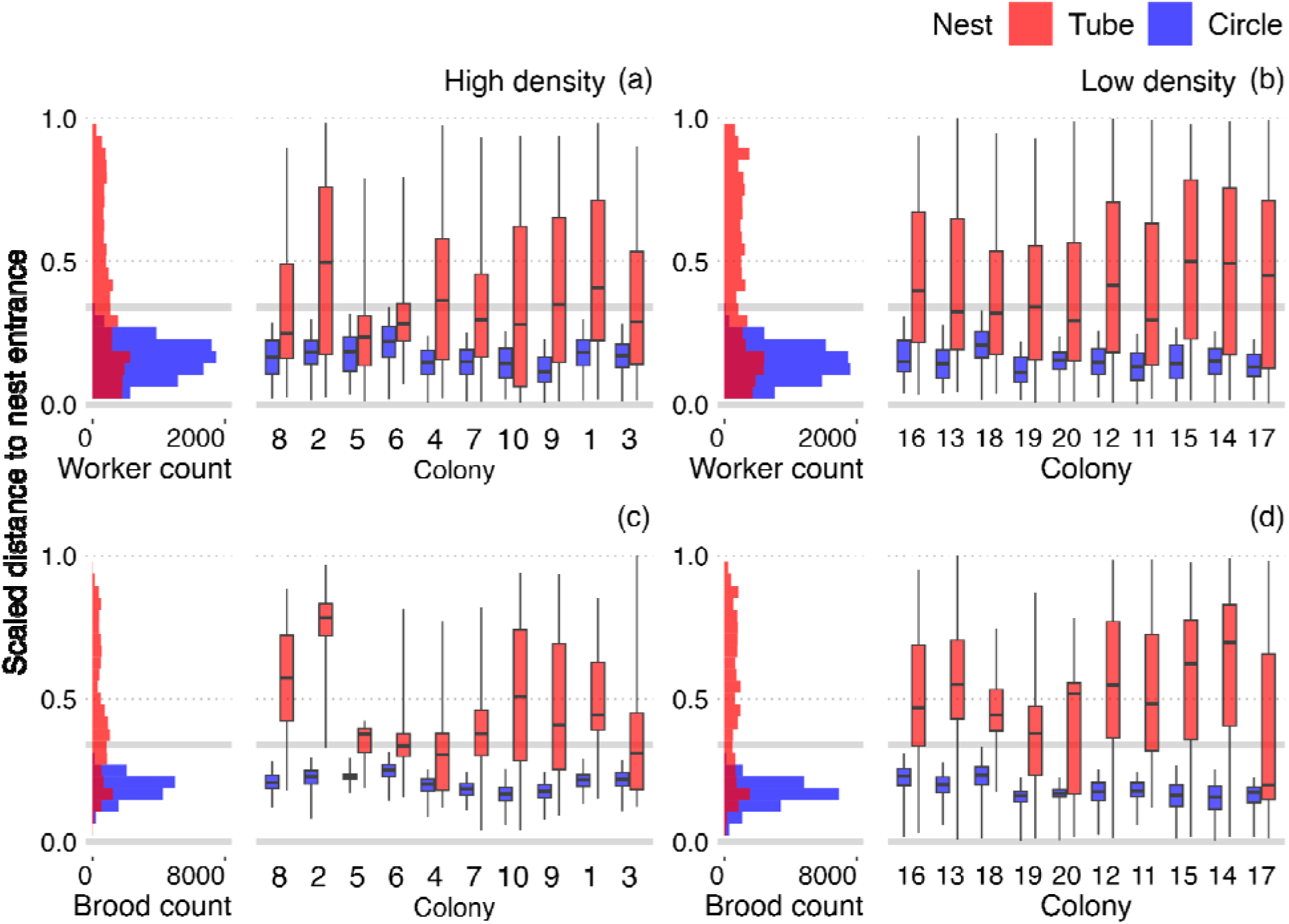
Actual distances of workers and brood from the nest entrance scaled only by nest size, by setting the farthest point of the tube nest as 1. When comparing these distances to the distributions across sections as in Fig. 3, it becomes clear how colonies in tube nests are spread out more from the entrance and from each other relative to the circle nest (where the range of farthest possible distances from the entrance is indicated by the gray lines). Shown are worker (a, b) and brood (c, d) distributions for the circle (blue) and tube (red) nests, and for high (a, c) and low (b, d) worker density treatments. N: workers = 30247 distance measurements, brood = 59459, queens = 1041, alates = 1006, over a total of 400 photos of 20 colonies. Colonies are sorted in order of increasing colony (and nest) size. For statistical analysis see Table A1.

### Distances from the center of the brood

As with distance to the entrance, we found that workers, queens, and alates were farther from the brood center in the tube nest (all p<0.01, Fig. 5, Table A2). We also found that male alates were found farther from the brood center than queen alates (Fig. A5d, Table A2), and that neither density nor time (‘Day’) had an effect (Table A2). Therefore, elongated nest shapes like our tube nest likely force colony members to real distances farther from unique points such as the nest entrance, the center of the brood, and the queen.

**Figure 5.**
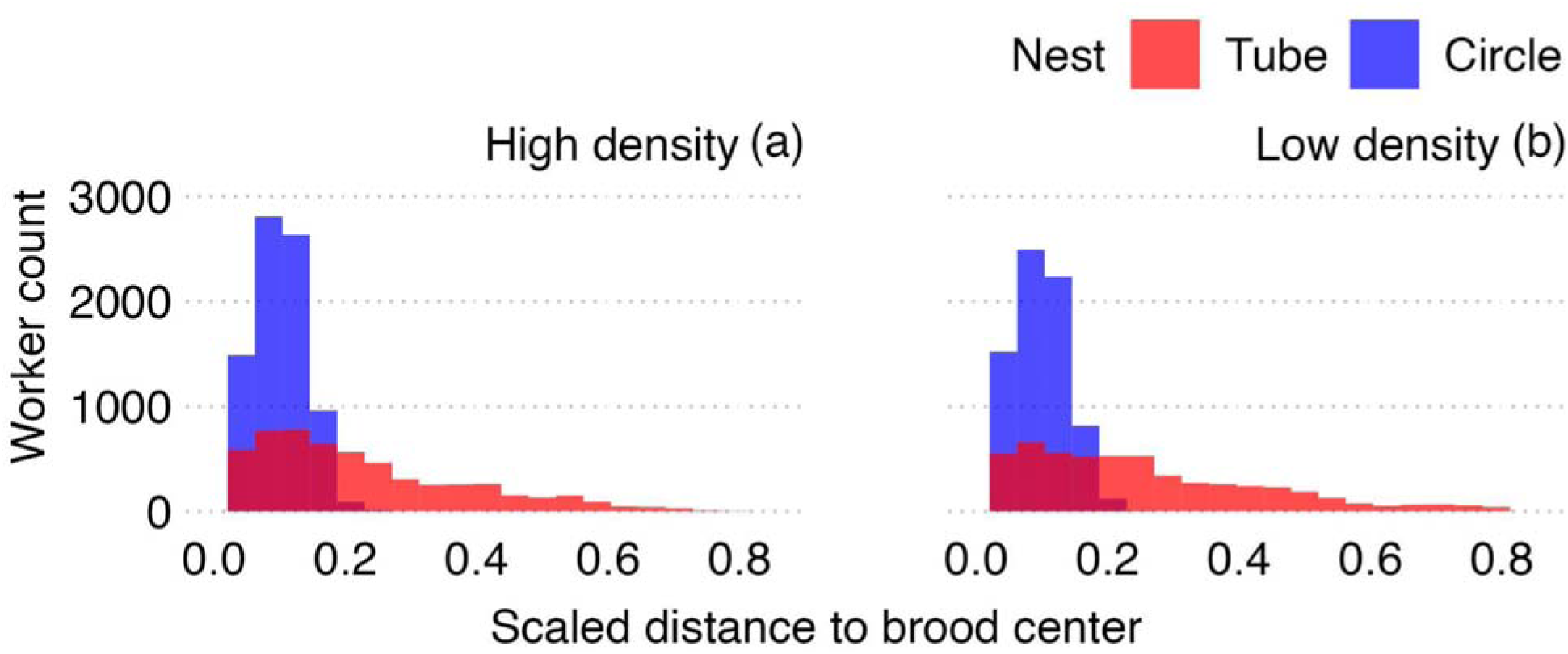
Workers are often dispersed farther from the center of the brood pile in the tube nest than what is possible in the circle nest (scaled only by colony size, by setting the farthest point of the tube nest as 1). Distributions are shown across high (a) and low (b) nest densities. N = 26733 observations of worker ants. For statistical analysis see Table A2.

### The effect of corners

The two nest shapes in our study were designed to maximally differ in their diameter/area ratio, i.e. elongation along a length axis, which simultaneously made one nest narrow, possibly limiting ants from assessing density in all parts of the nest. However, since we arranged both nests to be contained within our standard 2×3inch nest size, this also caused the tube nest to be ‘bent’, i.e. to form corners. The circle nest also has slight heterogeneities where the actual ‘circle’ is met by the short entrance tunnel. We found that these corners, which might have affected movement, affected spatial distributions of workers, brood, queens, and alates in the nests (with more workers, brood, and queens in ‘corners’, and fewer alates: all p<0.05 for the effect of ‘nest section including a corner’ on proportion of individuals in that section, Table 1).

### Small differences between colonies in spatial distributions

Colony identity was included as a random effect in the statistical models for both proximity to the nest entrance (Table A1) and to the center of the brood (Table A2). The impact of random effects can be estimated from the difference between marginal and conditional *R*^2^ values; here, we found that colony ID explained very little of the variation among worker distances (1.5% for the distance to the entrance, 2.2% for distances to the center of the brood), but more of the variation among for brood and queens (4.8% for distance of brood to entrance; 6.3% and 7.8% for of variation in distance to entrance and brood center, respectively, for queens), but a larger amount for the distances of alates from the entrance (17.2% and 0.6% respectively). This may reflect that the positions of alates are more idiosyncratic by colony and depend on the sex of the alates.

### Worker site fidelity is a function of colony size and density, not geometry…

Despite these significant impacts of nest geometry on the distances of colony members to each other and the distributions particularly of brood, we did not find any impact of nest shape on worker spatial fidelity (LME, p=0.96 for scaled zone sizes measured as number of nest sections, p=0.16 for actual zone sizes in cm^2^, N=384 worker spatial fidelity zones from a total of 81 unique workers, Fig. 6, Table 2) or occurrence (p=0.94 for scaled, p=0.10 for actual) zone sizes. Instead, the main predictor of how dispersed an individual’s movement through the nest were seemed to be the total nest size (which we scaled with colony size to keep a controlled worker density in our experiment; Fig. 7): colony size did not predict scaled zone sizes but did strongly correlate with absolute zone sizes, and also interacted significantly with density in determining absolute zone sizes (different slopes in Fig. 7 a vs b and c vs d). This (the strong effect of ‘Colony size’ but not density, and the effect on absolute zone area but not proportion of the nest used) is consistent with the idea that it is available area, not the number of individuals, that matters. The average distance of the worker’s location from the nest entrance or brood center impacted neither fidelity zone nor occurrence zone sizes (Tables 2 and A3, Fig. 6). Colonies differed only very slightly in their workers’ zone sizes (Colony ID explained 6-7% of variation in scaled zone sizes, and only 1% of variation in absolute zone sizes, Table 2).

**Figure 6.**
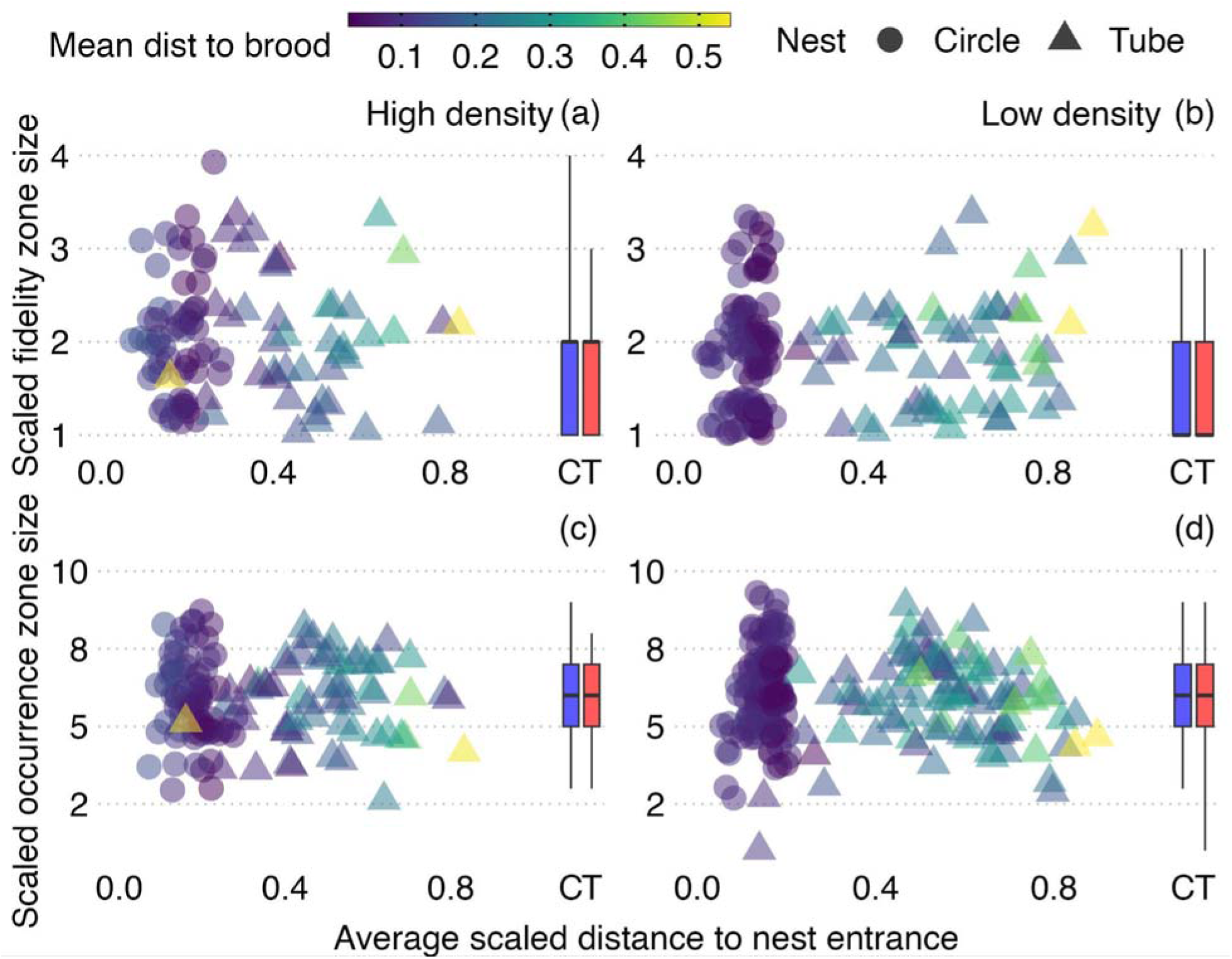
Worker spatial fidelity and occurrence zone sizes (in nest sections out of 24 total) were constant across the two nest shapes and did not differ with density, distance to the nest entrance (see Table 2 for statistics) or distance to the brood center (see Table A3 for statistics). Fidelity zones: sections in which workers spent at least 15% of observations: a, b; Occurrence occurrence zones: all sections in which a worker was observed: c, d. N: 383 marked ants across 19 colonies had sufficient observations to be included in the analysis (74 in circle/high density, 148 in circle/low density, 63 in tube/high density, 99 in tube/low density).

**Figure 7.**
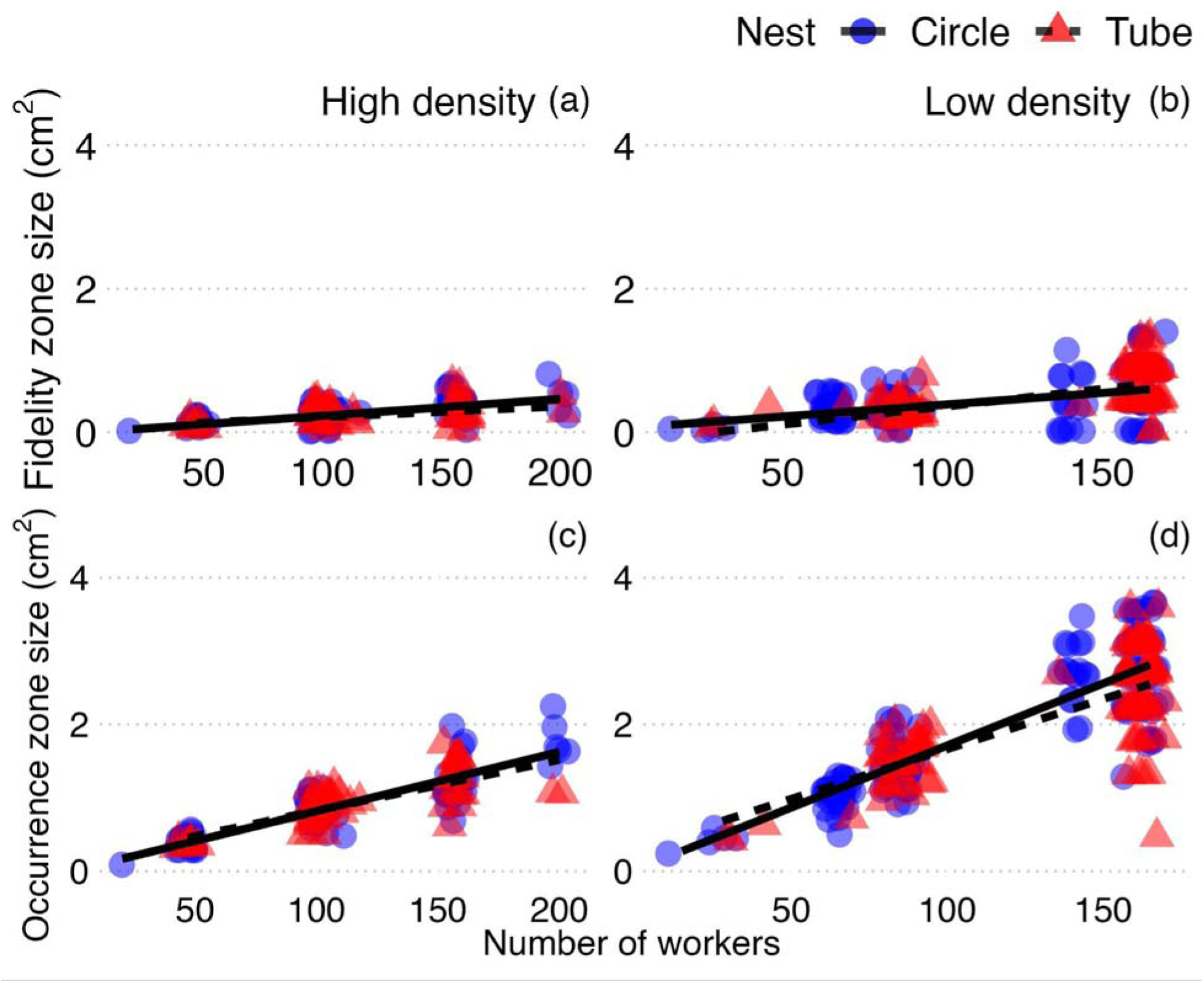
Worker spatial fidelity (a, b) and occurrence (c, d) absolute zone sizes (in cm^2^) increased with number of workers (i.e. colony size and thus total nest area, since all colonies were given nests of a size that was scaled to worker number). Nest shape (tube: red, circle: blue) and worker density (high: a, c; low: b, d) did not have significant main effects, although density interacted significantly with colony size. Only for occurrence zone size was there a significant interaction between colony size and nest shape (difference in slope in c/d). Scatter plot points are jittered by 5 x-axis units. Lines represent significant linear relationships (solid lines for circle nest, dashed lines for tube nest; Table 2). N: 384 ants as in Fig. 6.

**Table 2.**
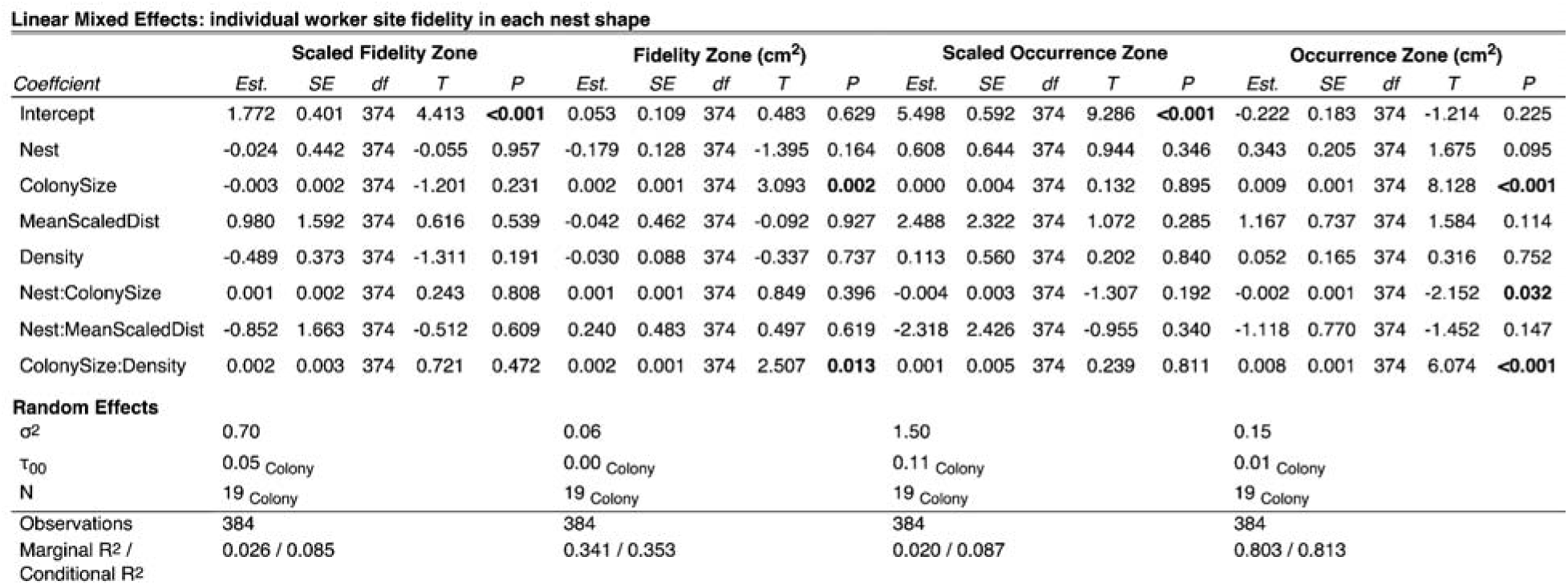
Size of worker spatial fidelity zones and occurrence zones related to nest shape (Nest), nest and colony size (ColonySize), density treatment (Density), and distance of the average position from the nest entrance (MeanScaledDist). Scaled zone sizes are as a proportion of total nest area, other zone sizes are in actual area. The random effect Colony ID is colony identity.

There was also a significant interaction effect of nest shape and colony size for the absolute size of occurrence zones (p=0.03); as seen in Fig. 7, occurrence zone sizes increase more slowly with the number of individuals (∼total available area) in the tube nest than the circle nest. Since the same was not found for the more conservatively defined fidelity zones (p=0.40), it is possible that this is because workers in the tube nest wandered into zones outside their typical fidelity zone more rarely in the tube nest, particularly in large nests. Bold *P* values indicate significance. Model formula in R: Zone ∼ Nest * ColonySize + Nest * MeanScaledDist + ColonySize * Density + (1 | ColonyID)

### …But geometry drives the location of spatial fidelity zones relative to points in the nest

Nest shape reverses the relationship between worker spatial fidelity zone locations and important points in the nest space. In the circle nest, workers who are close to the center of the brood pile are far from the entrance and vice versa; in the tube nest on the other hand, the opposite is the case: workers close to the entrance are also close to brood, and workers in the back of the nest are far from both the entrance and the center of the brood (LME, p<0.001 for interaction of nest shape and distance to the entrance on distance to the center of the brood, using the center of each worker’s spatial fidelity zone; Fig. 8, Table 3). Only a small portion of variation (3%, Table 3) was explained by the random effect of colony identity.

**Figure. 8.**
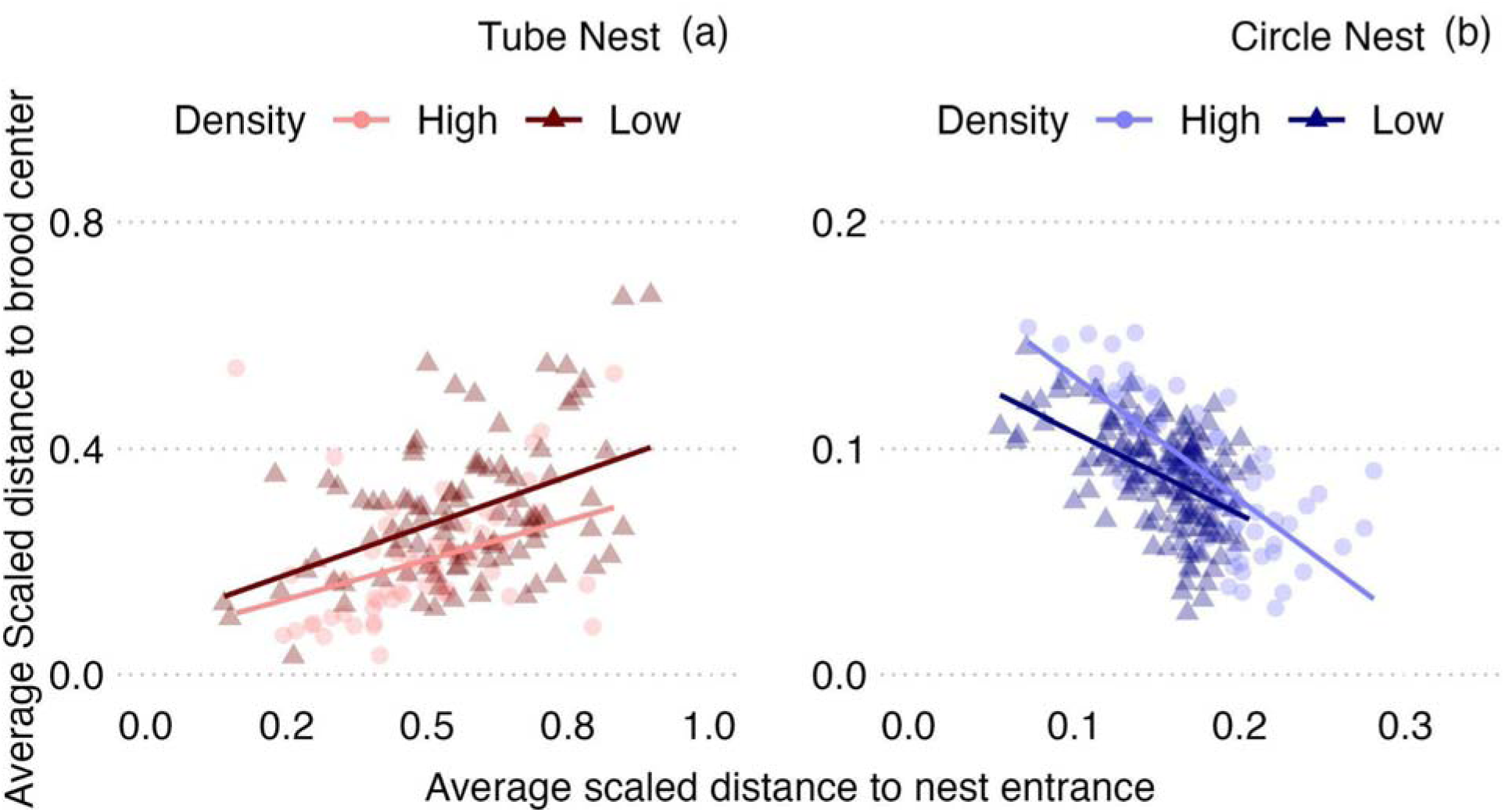
Worker spatial fidelity zones nearer the nest entrance are further away from the center of the brood in the circle nest (b), in agreement with previous literature using similar nest shapes; however, in tube-shaped nests (a), we find the opposite relationship: workers who are close to the entrance are also close to brood. Overall distances are longer for the circle nest (as it does not extend as far, see also Fig. 4). Worker density did not affect locations of spatial fidelity zones. For statistics, see Table 3.

**Table 3.**
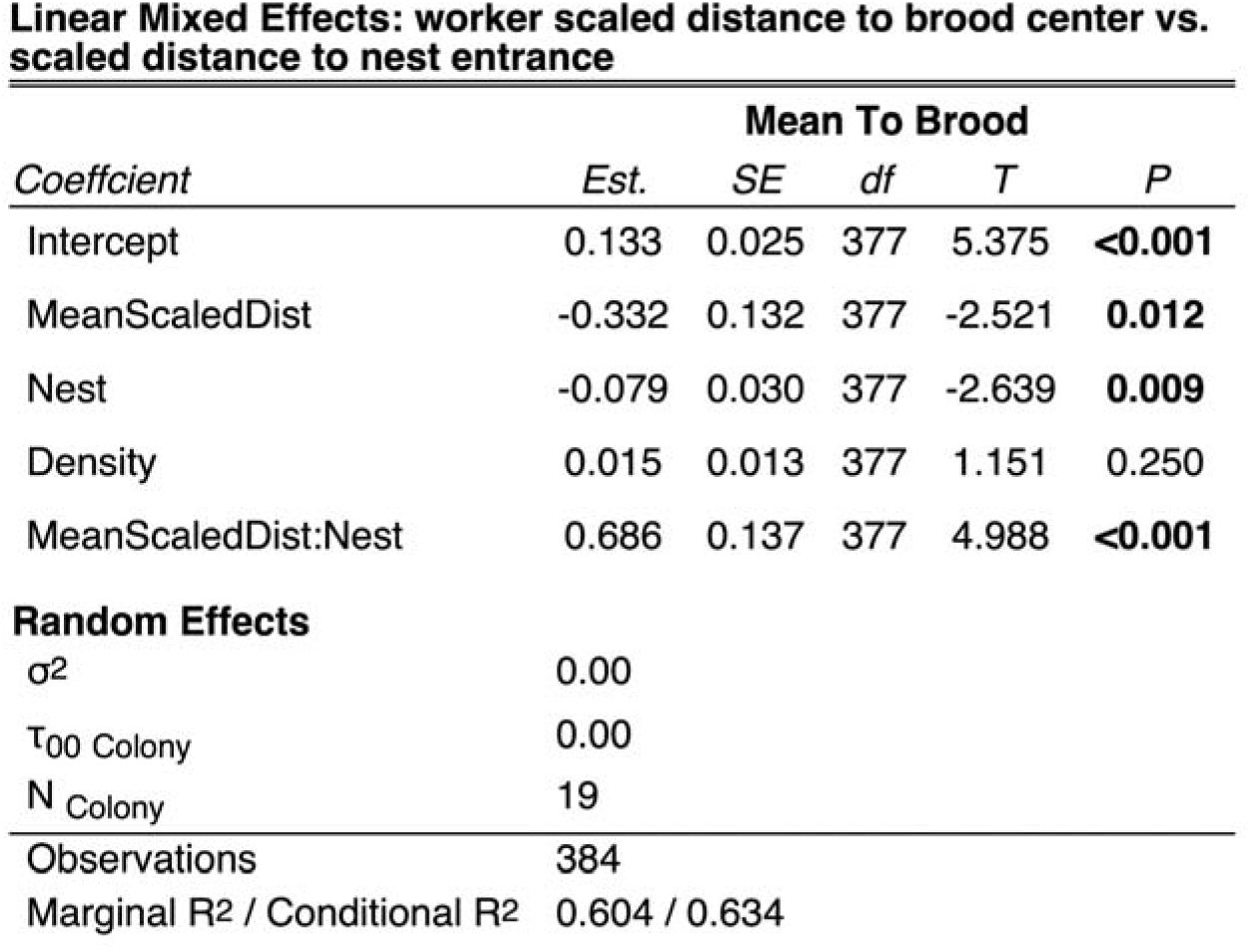
The relationship of worker spatial fidelity zone locations relative to the nest entrance and relative to the center of the brood (see also Fig. 8).

## Discussion

Available nest geometry influences the spatial organization of colonies in the rock-dwelling ant *Temnothorax rugatulus*. In our circle-shaped nest, as in previous studies using similar ‘open’ nest geometries, worker ants distributed themselves approximately evenly across the available nest space, while brood and queens were clustered at an intermediate distance from the entrance, and alates tended to be near the entrance. However, in the constricted space of the tube nest, brood were more evenly distributed, all types of colony members were at further distances from specific points (such as the nest entrance and the center of the brood), and workers, brood, and queen locations were affected by heterogeneities of the nest geometry such as corners. The tube geometry thus may have reduced global interaction, and may have prevented the clear distinction between nest areas with respect to function that is apparent in the circle nest (e.g. a distinct brood area separate from the space containing alates). Interestingly, overall worker density did not affect these patterns. In contrast to these pervasive effects of nest geometry on distributions and relative distances of colony members to each other, we found that individual worker spatial fidelity zone sizes remained constant across nest geometries. That is, the area of the nest that is used by each worker remains remarkably consistent as a proportion of total nest area across both nest shapes and nest sizes (scaled with colony size). In absolute terms, both conservative spatial fidelity zones (areas workers repeatedly visit) and general occurrence zones (all areas with observations for a specific worker) were largely determined by total available nest area (scaled with colony size, i.e. worker number), but did not change with nest geometry. One exception to this is that we found a significant interaction of colony size and nest shape in their effect on absolute occurrence zone (but not fidelity zone) sizes - this may indicate that at large nest and colony sizes, workers in circle nests make occasional forays into more different sections of the nests than workers in tube nests (even as their core spatial fidelity zones remain unchanged).

Does it matter how workers, brood, and queens are distributed across the space of a nest? Our results show that both nest size and nest geometry can affect parameters such as the clustering of colony features, the distances of individuals from key points, and heterogeneity in local density. Previous work has shown that in *Temnothorax* ants, nest area plays a key role in nest site selection (Pratt and Pierce 2001; Mitrus 2015), and workers actively gauge the size of prospective nests (Pratt 2005; Pavlic et al., 2021). It has also been demonstrated that worker density matters, as it can elevate worker energy expenditure (Cao and Dornhaus 2008) and lead to increased foraging and scouting rates and induce polydomy (Cao 2013). Additionally, with respect to geometry, certain structural features can alleviate consequences of high nest density like traffic jams during nest evacuation (Burd et al., 2010; Shiwakoti et al., 2014; Wang and Song 2016). While *Temnothorax* ants generally nest in preexising cavities, the size and geometry of which may constrain them, they can make some modifications, both by adding internal surrounding walls (Franks et al., 1992; Franks and Deneubourg 1997; Aleksiev et al., 2007a; Aleksiev et al., 2007b; Aleksiev et al., 2007c; DiRienzo and Dornhaus 2017), and because inactive colony members might act as temporary barriers, similar to how army ants produce walls to regulate trail traffic (Baudier and Pavlic, 2020). The purpose of such modifications is not yet clear, but it suggests that these ants might actively alter the available area or shape of their nests. We also know in general that ants may also regulate contact rates as part of strategies for task allocation and information exchange (Pacala et al., 1996; Pinter-Wollman et al., 2012; Pinter-Wollman et al., 2013; Lehue et al., 2020b), and that contact rates can be used to estimate colony or group sizes (Gordon 1992; Pratt 2005; Dornhaus and Franks 2006). Given that our study suggests that nest geometry may impact local density and distribution of workers relative to brood and queens, it is thus likely that both worker interaction rates and the interactions between workers and brood, and thus colony function generally, are also impacted. Future studies should closely examine how such interactions are affected by nest area and geometry, as well as possible heterogeneities in local densities and relative positions of different colony members.

While global spatial distributions and relationships across workers, brood, and queens were significantly affected by nest size and geometry, we found that nest shape did not affect either the size of workers’ spatial fidelity zones (a median of ∼4-8% of total nest area) nor the overall area used by each worker, which we termed ‘occurrence zones’ (∼25% of total nest area, i.e. approximately 6x the size of the fidelity zones for our method and sample size). We found that the absolute size of these zones (∼0.1 - 4 cm^2^ for fidelity zones) was primarily affected by the total nest size, which was artificially scaled with colony size. We can thus not definitively separate the effects of colony size (number of workers) and available nest area; however, since density had no main significant effect, it is likely that it is simply total nest area that determines spatial fidelity zone size. This is interesting since it indicates that workers somehow ‘divide up’ the available nest area among them, but not simply by aiming for a particular worker density. It is possible that workers move until achieving a constant density (interaction rate) over time. While our study provides these clues, it does not fully demonstrate the mechanism used by workers in determining either the placement or the size of their spatial fidelity zone.

The placement and size of worker spatial fidelity zones are thought to affect task allocation and specialization across workers, by virtue of determining which task items workers are likely to interact with, in *Temnothorax* (Franks and Tofts 1994; Leitner and Dornhaus 2019) and other social insects (Sendova-Franks and Franks 1995; Jandt and Dornhaus 2009; Powell and Tschinkel 1999). Given this potential importance of spatial fidelity zones for colony organization, it is thus possible that the ants in our study were actively regulating spatial fidelity zone size to match their needs independently of nest geometry. Such resilience could potentially be an evolved strategy to adapt to varying nest conditions, a hypothesis supported by the ecological diversity of *T. rugatulus* colonies across North America (Bengston and Dornhaus 2015) and the fact that in some populations, colonies face strong nest competition and thus constraints on nest selection (Bengston and Dornhaus 2015). In contrast to previous findings in the ant *Temnothorax unifasciatus* and the bumble bee *Bombus impatiens*, we did not find spatial fidelity zone size to change systematically across the nest space (e.g. larger zones further from the nest center, Sendova-Franks and Franks 1995; Jandt and Dornhaus 2009). However, our observations are in line with earlier studies on *T. rugatulus*, which showed that although inactive workers typically have small spatial fidelity zones and stay near the nest center, this pattern changes during their active phases (Charbonneau et al., 2017b).

Our study addresses the general principle of how external, spatial configuration can affect the behavior and function of adaptive systems. The importance of real space and physical constraints on ant colonies specifically (Pinter-Wollman et al., 2017) and more widely on self-organized and collective systems both engineered and biological (Araújo et al., 2023) is increasingly being recognized, and has wide-ranging implications (Araújo et al., 2023; Pinter-Wollman et al., 2017; Pinter-Wollman et al., 2018). Two processes are particularly important. First, confinement, i.e. in this case physical barriers, can affect self-organized pattern formation; and beyond just forcing a collective system to obey boundaries, can lead to entirely different outcomes with regard to relationships and functionality, even in counterintuitive ways (Schäfer et al., 2022). Second, stigmergy, i.e. the encoding of organizationally relevant information in the external environment, is a key principle that allows efficient and robust organization in many natural systems. In this study, we show that the brood clustering and the spatial separation of a dense brood area from other nest areas (e.g. the area near the entrance and area containing alates) disappears when ants are inhabiting a nest shape that, while offering the same total area, limits and defines traffic and interactions across nest sections. This, first, demonstrates the role that confinement can play in structuring or destroying self-organized patterns. Second, if, as hypothesized for these ants, task specialization is organized by worker spatial fidelity zones overlapping with only specific task stimuli, this stigmergic organization is upended by different nest shapes that prevent ants from separating out different task areas. Both confinement and stigmergy thus drive important aspects of self-organized function and we demonstrate here that they can be significantly impacted by external spatial constraints.

## Appendix

Table A1. Relationships between colony members scaled distance to the nest entrance and nest shape and physical properties. Sex is male or female alates, and ratio is the proportion of male alates in the observation. Formula: ScaledDist ∼ Nest*Density + Day + (1 | Colony ID); Formula (alates): Formula: ScaledDist ∼ Nest + Day + Sex + SexRatio (1 | Colony ID)

Table A2. Relationships between mobile colony members scaled distance to the brood center and nest shape and physical properties. The brood center is a colony’s brood centroid in each observation. Distance to the brood center is the absolute value of the difference between the scaled distances of the brood center from each worker to the entrance and is scaled such that 1 is the back of the tube nest. Formula: ToBrood∼ Nest*Density + Day + (1 | Colony ID); Formula (alates): Formula: ToBrood ∼ Nest + Day + Sex + SexRatio (1 | Colony ID)

Table A3. The relationship between worker site fidelity and scaled distance to the brood center. Scaled fidelity zone size is the summation of all zones that a worker was found in for at least 15% of its observations, and scaled occurrence zone size counts all zones with any observations (and divide by the total of 24 possible, equal-area zones; worker must have at least 7 observations). True zone size (cm^2^) multiplies this number by the internal area of the nest (note that nest size was scaled to worker number and thus differs between colonies). Formula: Zone ∼ Nest * ColonySize + Nest * MeanToBrood + Density * ColonySize + (1 | Colony)

Figure A1. The reference photo used to determine the nest area allocated to each worker in a colony. The colony contains 248 individuals. The internal area is approximately 4.11mm^2^, producing 0.017 mm^2^ for each worker, which was doubled to 0.033 mm^2^ to promote more flexible space usage. This value represents the high nest density treatment and was doubled to produce the low nest density treatment: 0.066 mm^2^ for each worker in a colony.

Figure A2. The eight equal-area nest sections for the circle (a) and tube (b) nests (black lines) used to determine the densities of colony members through the nest, and the twenty-four equal area zones used to calculate worker site fidelity in the circle (c) and tube (d) nests. Two zone sizes were used because the larger sections potentially capture tasks throughout the nest whereas the smaller sections capture the smaller grain worker spatial fidelity zones.

Figure A3. Visual example of the shortest distance from an example worker (red dots) to an example brood center (white dots) in the circle (a) and tube (b) nests. We found each colony member’s distance to the nest section closest to the entrance using reference coordinates for that nest section, where we then added the shortest distance from that nest section to the entrance through reference coordinates. In the circle nest, distance to the entrance = sqrt((colony member x - entrance x)^2^ + (colony member y - entrance y)^2^), in the tube nest, where direct (i.e. straight-line) path was often not possible within the available nest space, distance to the entrance = sqrt((colony member x - nest section entrance x)^2^ + (colony member y - nest section entrance y)^2^) + reference distance, where the nest section entrance is the closest accessible facing toward the direction of the entrance (Fig. A3b). Correcting corner cuts at the entrance: Near the entrance of both nests (in nest section 1), a colony member’s distance to the nest entrance can cut through the corner where the entrance tunnel opens into the nest (Fig. A3c). We solved this by first determining whether a colony member’s distance would cut through the corner and then assigning an alternative distance to the entrance where necessary.

Subfigure (c): the criteria used to determine if a colony member required an alternative linear distance to the nest entrance near the corner formed from the nest entrance tunnel opening into the nest (black solid lines). The red dot represents a colony member, and the black dot represents the entrance. The dashed lines represent the hypotenuses between the colony member and the corner (red) and the corner and the entrance (black). The arcs represent angles, and the black square represents 90°. The solid red polygon indicates nest space that would require this type of alternative calculation of distance to the nest entrance.

Figure A4. When showing the actual distances of queens (a, b) and alates (c, d) from the nest entrance (scaled only by colony size, by setting the farthest point of the tube nest as a distance of 1) instead of sections as in Fig. 3, it becomes clear how colony reproductive members in tube nests (red) are spread out more from the entrance and from each other relative to the circle nest (blue). This observation holds true regardless of high (a) and low (b) nest densities. Alate males are also found to be closer to the nest entrance than queen alates (d). Sample sizes were 1178 queen observations across 20 colonies, and 1006 alate observations across 7 low density treatment colonies.

Figure A5. Both queens and alates are found farther from the center of the brood pile (centroid) in the tube nest (red) than what is possible in the circle nest (blue) (scaled only by colony size, by setting the farthest point of the tube nest as 1). Differences in queen distributions are shown across high (a) and low (b) nest densities; alates were only present in the low-density treatment. Alate males are found to be farther from the brood centroid than young queens (female alates, d). Sample sizes were 1080 queen observations from 400 photos across 20 colonies in each density treatment, and 676 alate observations across 7 low density treatment colonies.

