## Supplementary figures and images for "Cavity geometry shapes overall ant colony organization through spatial limits but workers maintain fidelity zones"

### Figure A1

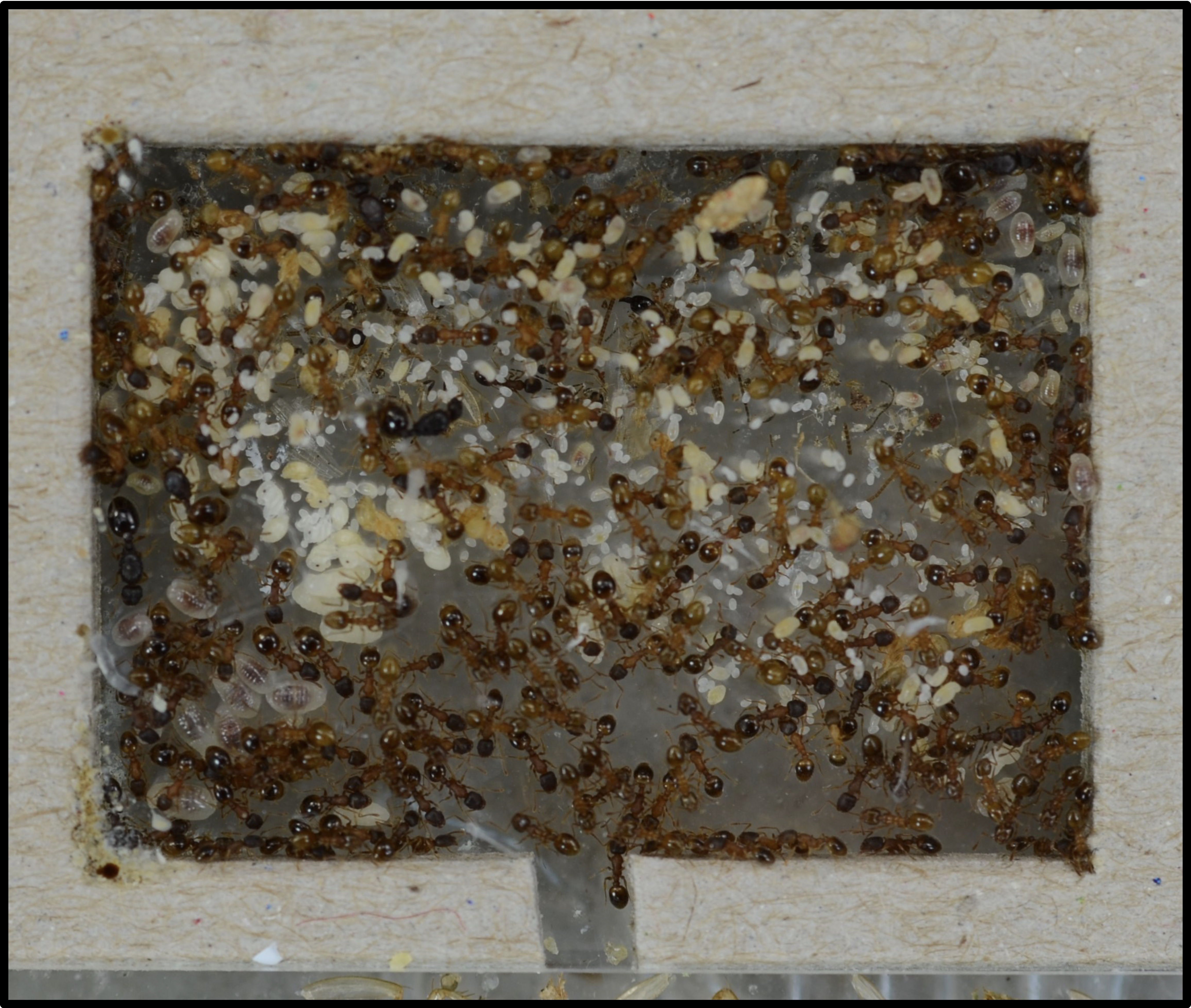

### Figure A2

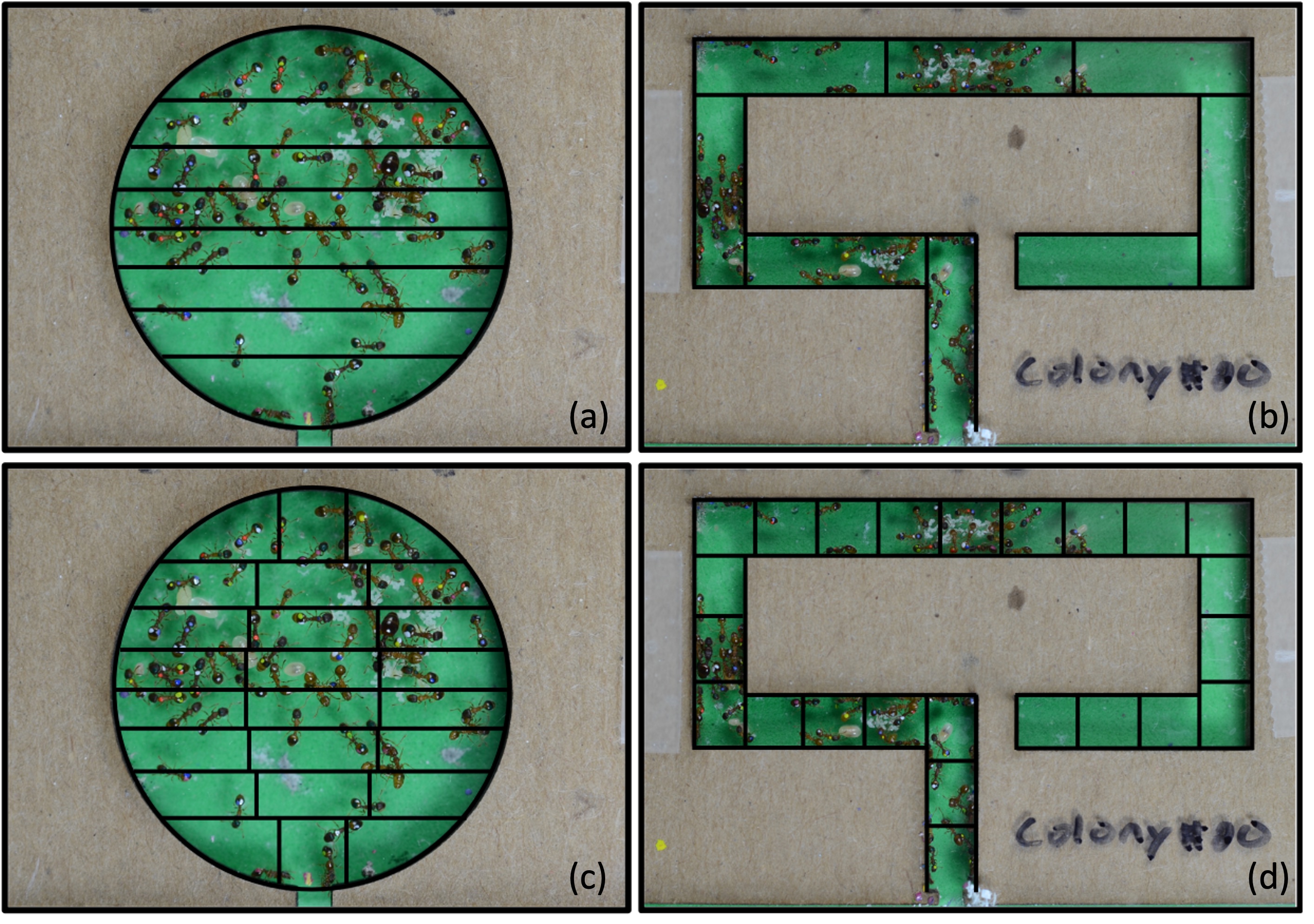

### Figure A3

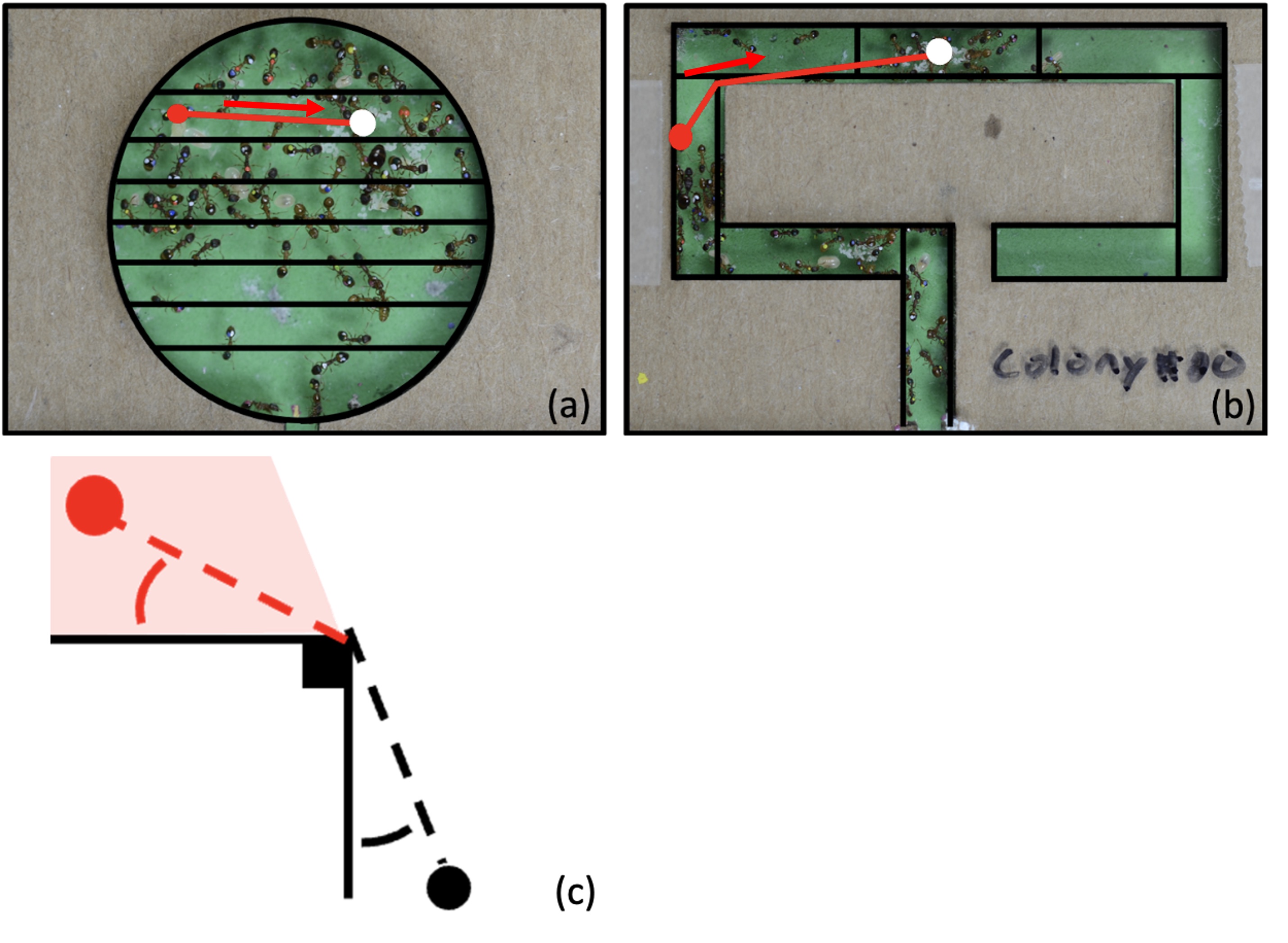

### Figure A4

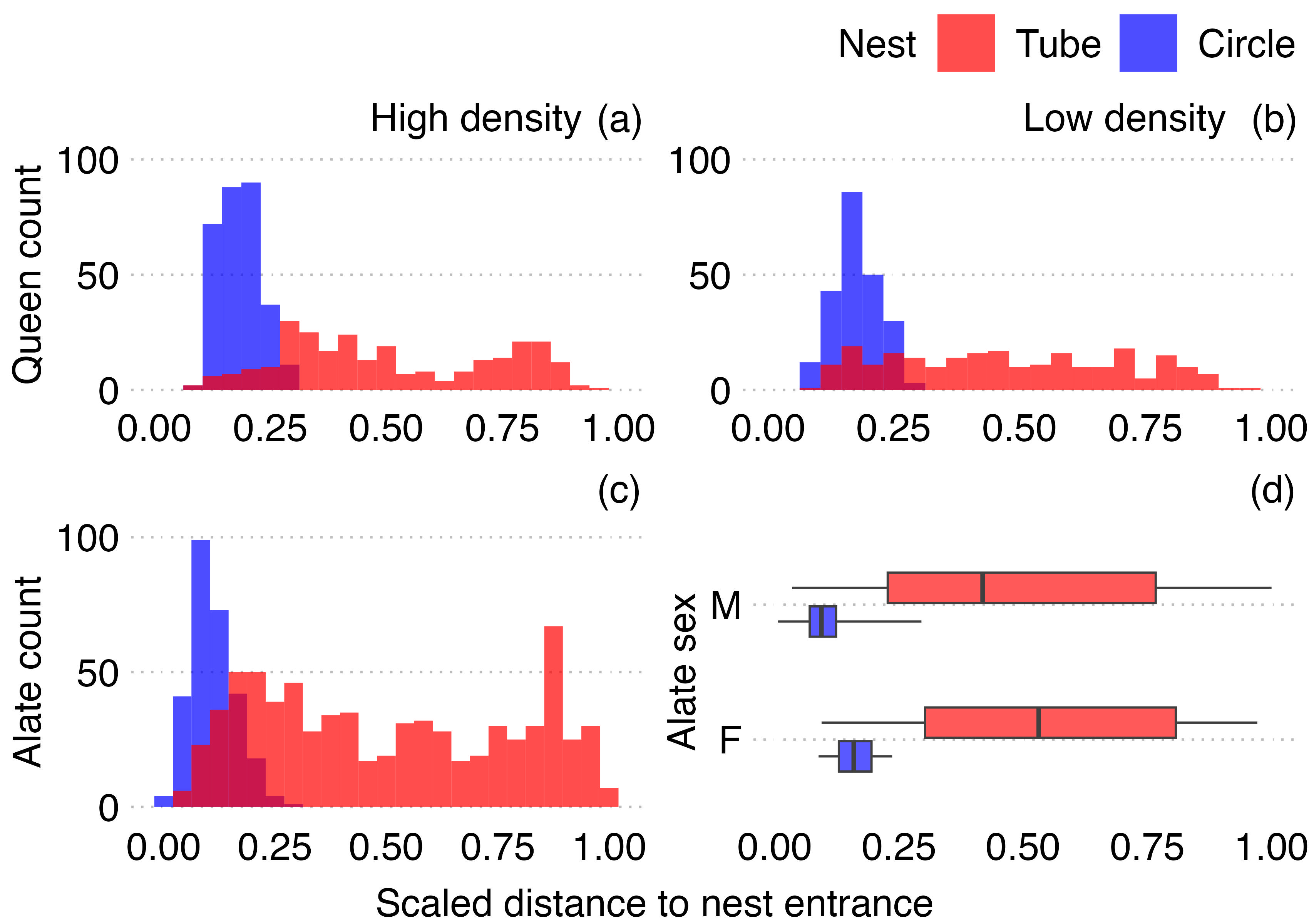

### Figure A5

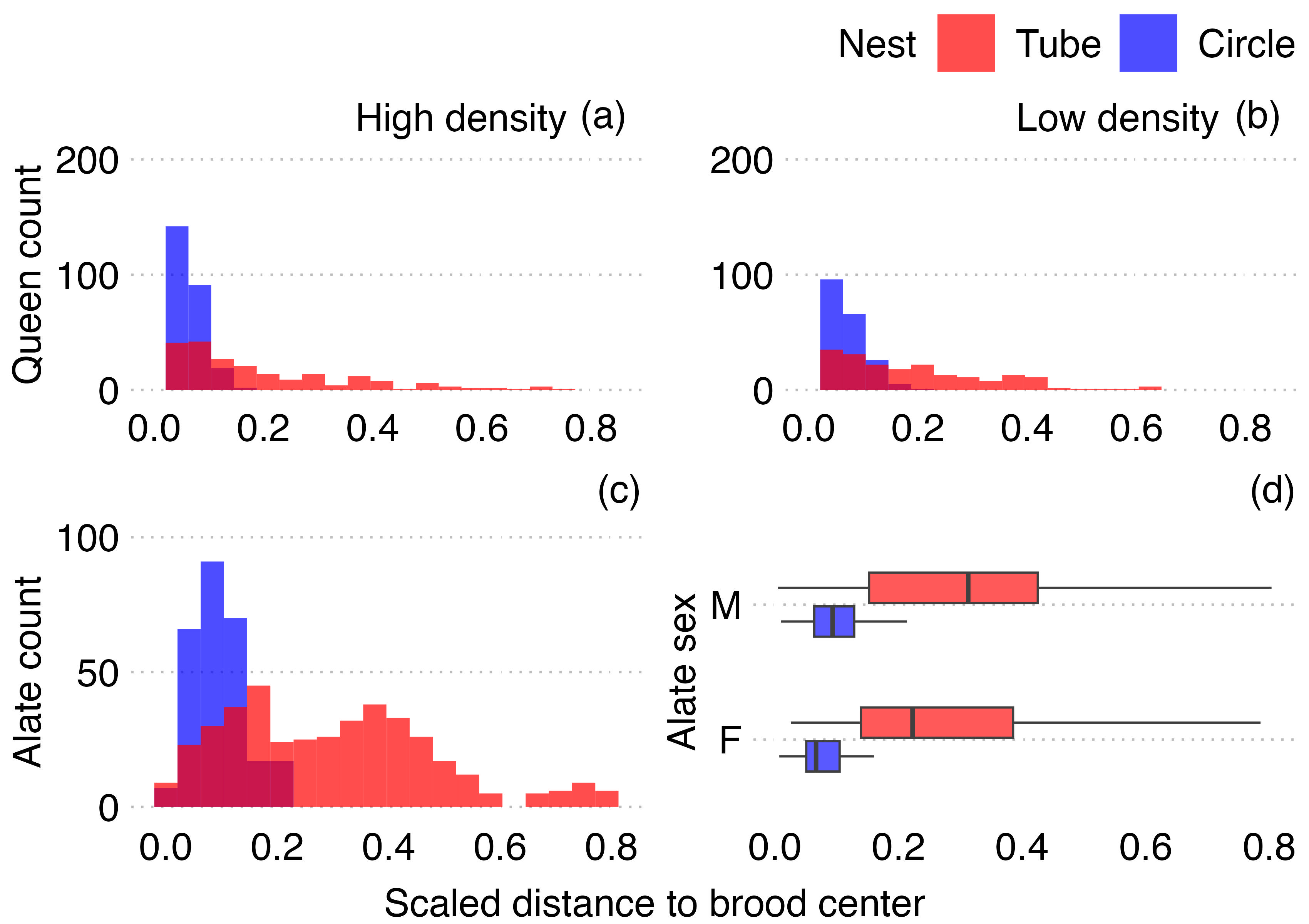
