## Supplementary material for "Cavity geometry shapes overall ant colony organization through spatial limits but workers maintain fidelity zones": Table A1

**Linear Mixed Effects: colony member scaled distances to the nest entrance**

| | Workers | | | | | Brood | | | | | Queens | | | | | Alates | | | | |
|---|---|---|---|---|---|---|---|---|---|---|---|---|---|---|---|---|---|---|---|---|
| *Coeffcient* | *Est.* | *SE* | *df* | *T* | *P* | *Est.* | *SE* | *df* | *T* | *P* | *Est.* | *SE* | *df* | *T* | *P* | *Est.* | *SE* | *df* | *T* | *P* |
| Intercept | 0.150 | 0.010 | 30240 | 15.407 | **<0.001** | 0.193 | 0.017 | 59452 | 11.635 | **<0.001** | 0.195 | 0.025 | 1034 | 7.966 | **<0.001** | 0.297 | 0.090 | 999 | 3.312 | **0.001** |
| Nest | 0.230 | 0.003 | 30240 | 70.377 | **<0.001** | 0.258 | 0.002 | 59452 | 124.010 | **<0.001** | 0.324 | 0.013 | 1034 | 24.892 | **<0.001** | 0.559 | 0.044 | 999 | 12.688 | **<0.001** |
| Density | -0.021 | 0.013 | 30240 | -1.590 | 0.112 | -0.028 | 0.023 | 59452 | -1.213 | 0.225 | 0.001 | 0.031 | 1034 | 0.022 | 0.982 | | | | | |
| Day | 0.001 | 0.000 | 30240 | 4.409 | **<0.001** | 0.001 | 0.000 | 59452 | 3.338 | **0.001** | -0.001 | 0.001 | 1034 | -1.106 | 0.269 | -0.009 | 0.003 | 999 | -3.515 | **<0.001** |
| Nest:Density | 0.061 | 0.004 | 30240 | 13.488 | **<0.001** | 0.072 | 0.003 | 59452 | 25.861 | **<0.001** | -0.022 | 0.019 | 1034 | -1.141 | 0.254 | | | | | |
| Sex | | | | | | | | | | | | | | | | -0.065 | 0.019 | 999 | -3.428 | **0.001** |
| SexRatio | | | | | | | | | | | | | | | | -0.080 | 0.072 | 999 | -1.119 | 0.263 |
| **Random Effects** | | | | | | | | | | | | | | | | | | | | |
| $\sigma^2$ | 0.04 | | | | | 0.03 | | | | | 0.02 | | | | | 0.05 | | | | |
| $\tau_{00}$ | $0.00_{\text{Colony}}$ | | | | | $0.00_{\text{Colony}}$ | | | | | $0.00_{\text{Colony}}$ | | | | | $0.03_{\text{Colony}}$ | | | | |
| N | $20_{\text{Colony}}$ | | | | | $20_{\text{Colony}}$ | | | | | $20_{\text{Colony}}$ | | | | | $7_{\text{Colony}}$ | | | | |
| Observations | 30247 | | | | | 59459 | | | | | 1041 | | | | | 1006 | | | | |
| Marginal $R^2$ / Conditional $R^2$ | 0.317 / 0.332 | | | | | 0.433 / 0.484 | | | | | 0.479 / 0.547 | | | | | 0.449 / 0.627 | | | | |
