## Supplementary material for "Cavity geometry shapes overall ant colony organization through spatial limits but workers maintain fidelity zones": Table A2

**Linear Mixed Effects: colony member scaled distances to the brood center**

| | Workers | | | | | Queens | | | | | Alates | | | | |
|---|---|---|---|---|---|---|---|---|---|---|---|---|---|---|---|
| *Coeffcient* | *Est.* | *SE* | *df* | *T* | *P* | *Est.* | *SE* | *df* | *T* | *P* | *Est.* | *SE* | *df* | *T* | *P* |
| Intercept | 0.098 | 0.007 | 26726 | 14.448 | **<0.001** | 0.054 | 0.016 | 941 | 3.420 | **0.001** | 0.100 | 0.039 | 669 | 2.582 | **0.010** |
| Nest | 0.125 | 0.002 | 26726 | 61.379 | **<0.001** | 0.127 | 0.009 | 941 | 13.776 | **<0.001** | 0.216 | 0.017 | 669 | 12.417 | **<0.001** |
| Density | -0.009 | 0.009 | 26726 | -0.951 | 0.341 | 0.028 | 0.019 | 941 | 1.522 | 0.128 | | | | | |
| Day | 0.000 | 0.000 | 26726 | 1.119 | 0.263 | -0.002 | 0.001 | 941 | -1.739 | 0.082 | -0.004 | 0.002 | 669 | -1.957 | 0.051 |
| Nest:Density | 0.030 | 0.003 | 26726 | 10.488 | **<0.001** | -0.016 | 0.014 | 941 | -1.187 | 0.236 | | | | | |
| Sex | | | | | | | | | | | 0.044 | 0.015 | 669 | 2.939 | **0.003** |
| SexRatio | | | | | | | | | | | -0.017 | 0.043 | 669 | -0.396 | 0.692 |
| **Random Effects** | | | | | | | | | | | | | | | |
| $\sigma^2$ | 0.01 | | | | | 0.01 | | | | | 0.02 | | | | |
| $\tau_{00}$ | 0.00 $_{Colony}$ | | | | | 0.00 $_{Colony}$ | | | | | 0.00 $_{Colony}$ | | | | |
| N | 20 $_{Colony}$ | | | | | 20 $_{Colony}$ | | | | | 7 $_{Colony}$ | | | | |
| Observations | 26733 | | | | | 948 | | | | | 676 | | | | |
| Marginal $R^2$ / Conditional $R^2$ | 0.268 / 0.291 | | | | | 0.251 / 0.327 | | | | | 0.339 / 0.344 | | | | |
