## Supplementary material for "Cavity geometry shapes overall ant colony organization through spatial limits but workers maintain fidelity zones": Table A3

**Linear Mixed Effects: individual worker site fidelity v. scaled distance to the brood center**

| | Scaled Fidelity Zone | | | | | Fidelity Zone ($cm^2$) | | | | | Scaled Occurrence Zone | | | | | Occurrence Zone ($cm^2$) | | | | |
|---|---|---|---|---|---|---|---|---|---|---|---|---|---|---|---|---|---|---|---|---|
| *Coeffcient* | *Est.* | *SE* | *df* | *T* | *P* | *Est.* | *SE* | *df* | *T* | *P* | *Est.* | *SE* | *df* | *T* | *P* | *Est.* | *SE* | *df* | *T* | *P* |
| Intercept | 0.105 | 0.093 | 374 | 1.131 | 0.259 | 0.105 | 0.093 | 374 | 1.131 | 0.259 | -0.004 | 0.162 | 374 | -0.026 | 0.979 | -0.004 | 0.162 | 374 | -0.026 | 0.979 |
| Nest | -0.185 | 0.104 | 374 | -1.777 | 0.076 | -0.185 | 0.104 | 374 | -1.777 | 0.076 | 0.089 | 0.173 | 374 | 0.518 | 0.605 | 0.089 | 0.173 | 374 | 0.518 | 0.605 |
| ColonySize | 0.002 | 0.001 | 374 | 3.215 | **0.001** | 0.002 | 0.001 | 374 | 3.215 | **0.001** | 0.009 | 0.001 | 374 | 8.042 | **<0.001** | 0.009 | 0.001 | 374 | 8.042 | **<0.001** |
| MeanToBrood | -0.650 | 0.660 | 374 | -0.986 | 0.325 | -0.650 | 0.660 | 374 | -0.986 | 0.325 | -0.071 | 1.062 | 374 | -0.067 | 0.947 | -0.071 | 1.062 | 374 | -0.067 | 0.947 |
| Density | -0.029 | 0.083 | 374 | -0.353 | 0.724 | -0.029 | 0.083 | 374 | -0.353 | 0.724 | 0.025 | 0.165 | 374 | 0.150 | 0.881 | 0.025 | 0.165 | 374 | 0.150 | 0.881 |
| Nest:ColonySize | 0.001 | 0.001 | 374 | 0.906 | 0.366 | 0.001 | 0.001 | 374 | 0.906 | 0.366 | -0.002 | 0.001 | 374 | -2.305 | **0.022** | -0.002 | 0.001 | 374 | -2.305 | **0.022** |
| Nest:MeanToBrood | 0.875 | 0.681 | 374 | 1.286 | 0.199 | 0.875 | 0.681 | 374 | 1.286 | 0.199 | 0.493 | 1.095 | 374 | 0.450 | 0.653 | 0.493 | 1.095 | 374 | 0.450 | 0.653 |
| ColonySize:Density | 0.002 | 0.001 | 374 | 2.619 | **0.009** | 0.002 | 0.001 | 374 | 2.619 | **0.009** | 0.008 | 0.001 | 374 | 6.039 | **<0.001** | 0.008 | 0.001 | 374 | 6.039 | **<0.001** |
| **Random Effects** | | | | | | | | | | | | | | | | | | | | |
| $\sigma^2$ | 0.06 | | | | | 0.06 | | | | | 0.15 | | | | | 0.15 | | | | |
| $\tau_{00}$ | 0.00 $_{Colony}$ | | | | | 0.00 $_{Colony}$ | | | | | 0.01 $_{Colony}$ | | | | | 0.01 $_{Colony}$ | | | | |
| N | 19 $_{Colony}$ | | | | | 19 $_{Colony}$ | | | | | 19 $_{Colony}$ | | | | | 19 $_{Colony}$ | | | | |
| Observations | 384 | | | | | 384 | | | | | 384 | | | | | 384 | | | | |
| Marginal $R^2$ / Conditional $R^2$ | 0.345 / 0.352 | | | | | 0.345 / 0.352 | | | | | 0.803 / 0.813 | | | | | 0.803 / 0.813 | | | | |
